# Neuro-computational account of arbitration between imitation and emulation during human observational learning

**DOI:** 10.1101/828723

**Authors:** Caroline C. Charpentier, Kiyohito Iigaya, John P. O’Doherty

## Abstract

In observational learning (OL), organisms learn from observing the behavior of others. There are at least two distinct strategies for OL. Imitation involves learning to repeat the previous actions of other agents, while in emulation, learning proceeds from inferring the goals and intentions of others. While putative neural correlates for these forms of learning have been identified, a fundamental question remains unaddressed: how does the brain decides which strategy to use in a given situation? Here we developed a novel computational model in which arbitration between the strategies is determined by the predictive reliability, such that control over behavior is adaptively weighted toward the strategy with the most reliable prediction. To test the theory, we designed a novel behavioral task in which our experimental manipulations produced dissociable effects on the reliability of the two strategies. Participants performed this task while undergoing fMRI in two independent studies (the second a pre-registered replication of the first). Behavior manifested patterns consistent with both emulation and imitation and flexibly changed between the two strategies as expected from the theory. Computational modelling revealed that behavior was best described by an arbitration model, in which the reliability of the emulation strategy determined the relative weights allocated to behavior for each strategy. Emulation reliability - the model’s arbitration signal - was encoded in the ventrolateral prefrontal cortex, temporoparietal junction and rostral cingulate cortex. Being replicated across two fMRI studies, these findings suggest a neuro-computational mechanism for allocating control between emulation and imitation during observational learning.

## Introduction

Learning from observing others is key to any social species. Whether it is learning a new skill by observing an expert or parent perform it, learning to seek positive outcomes or avoid negative outcomes, or making complex strategic decisions, observational learning (OL) is prevalent in our daily lives and presents adaptive advantages over experiential learning. Indeed, any species able to learn the consequences of actions available in their environment from observing their conspecifics possesses an evolutionary advantage, as it endows an individual with the ability to learn about the world without exposing oneself to the risks associated with having to directly sample those actions.

Two distinct strategies for observational learning have been proposed ^1–3^: action imitation and emulation. In action imitation, individuals learn by repeating actions that were most frequently performed by the other agent in the recent past, and in emulation learning, individuals learn by inferring the other agent’s goals, beliefs, intentions and/or hidden mental states (see ^4,5^ for reviews). Computationally, action imitation has been proposed to be accounted for in a reinforcement learning framework, whereby the actions performed by the other agent are reinforced via the computation of an action prediction error (APE) – the difference between the other agent’s action and how expected this action was. Evidence for a representation of APEs in the brain during imitation learning was found in dorsomedial and dorsolateral prefrontal cortex (dmPFC, dlPFC) and inferior parietal cortex ^6,7^, with the hypothesis that action imitation could be implemented in part through involvement of the mirror neuron system, active both when an action is observed and performed ^8–12^. In contrast, emulation learning consists of a more complex and flexible inference process. Several computational accounts of emulation have been provided ^13–15^, which can be collectively approximated as a form of Bayesian inference, whereby prior beliefs about the other agents are computed and combined with the evidence received from observation to produce posterior updated beliefs. These inference processes were found to recruit regions of the mentalizing network ^16^, specifically dmPFC, temporoparietal junction (TPJ) and posterior superior temporal sulcus (pSTS) ^15,17–21^.

However, if these two distinct OL strategies exist alongside each other in the brain, a fundamental open question remains: how does the brain decide which of these two strategies should be deployed in a given situation, and under what conditions does one or other strategy guide behavior? One possibility is that both strategies are simultaneously deployed and that behavior is a constant, static mix of the two. Alternatively, the brain might deploy an arbitration process whereby the influence over behavior of these strategies is dynamically modulated as a function of which strategy is deemed most suitable to guide behavior at a particular moment in time. The goal of the present study is to develop a computational model of arbitration between the two OL strategies, and to test this model against both behavioral and neural data obtained from human participants in order to understand how control is allocated between emulation and imitation strategies at behavioral, computational and neural levels.

In the domain of experiential learning, there is clear evidence that behavior is controlled by multiple competing systems, such as habits versus goal-directed actions ^22,23^ or model-free (MF) versus model-based (MB) learning ^24,25^. In order to allocate control to each system over behavior and to ensure control flexibly adapts to changes the environment, an arbitration mechanism has been proposed ^26^. A specific computational implementation of such an arbitration mechanism was suggested for MB and MF experiential learning ^27^, in which the reliability of the predictions of the two systems is dynamically computed by leveraging prediction errors generated from the two systems. In the brain, the ventrolateral prefrontal cortex (vlPFC) and frontopolar cortex (FPC) were found to encode the output of a comparison between the reliability of MF learning and MB learning. This suggests that in the experiential domain, depending on which learning strategy is more reliable at a given time point, the brain can allocate control over behavior to the most reliable system.

Whether a similar arbitration mechanism exists in the domain of observational learning remains unknown. Using similar principles to the reliability-mediated arbitration found in the experiential domain, we hypothesized that the allocation of control in OL between emulation and imitation strategies might also be related to the degree of uncertainty inherent in the predictions of the two systems. We therefore constructed a computational model of OL arbitration that utilized the degree of uncertainty in the predictions of the imitation and emulation systems in order to allocate behavioral control over these systems, so that across time, one or other system can come to dominate control over behavior depending on the relative uncertainties in the two system’s predictions. Concretely, given that the imitation system is aiming to keep track of predictions about which actions an observed agent will perform, we hypothesized that if the observed agent’s actions were to become more stochastic from trial-to-trial, the uncertainty in the imitation model predictions would decrease, thereby resulting in emulation becoming more favored. Conversely, we hypothesized that if the goal inference process were to become more uncertain because of task conditions (and also more difficult), then the predictions of the emulation system would become more uncertain, thereby favoring the imitation system.

With these strong theoretical motivations in mind, we designed a completely novel observational learning task (**Fig. 1**), in which we induced changes in the experimental conditions in order to maximize differential engagement of the two strategies depending on the environment, as hypothesized by our model. Two groups of 30 participants, referred to as Study 1 (initial sample) and Study 2 (replication sample) completed the task while undergoing fMRI scanning. The methods, computational modelling, behavioral analyses, fMRI pipeline, and results of Study 1 were pre-registered on OSF (project: https://osf.io/49ws3/; pre-registration: https://osf.io/37xyq) before Study 2 data collection. The use of pre-registration in this context allowed us to evaluate the extent to which our findings are susceptible to potential concerns about considerable modeler and experimenter degrees of freedom that can exist in either computational modeling and neuroimaging studies. The analytical flexibility that can apply in both types of studies has been suggested to lead to potential pitfalls of overfitting, thereby compromising the generalizability and reproducibility of findings ^28–30^. Here, by indulging in pre-registration of our computational models, and our full model-fitting procedures, fMRI pre-processing and fMRI statistical analysis pipeline, we reduce both modeler and experimenter degrees of freedom markedly. In our 2^nd^ experiment, we are able to report a full out-of-sample validation of both our computational modeling findings on behavior as well as on neuroimaging results. Thus, this study has important implications for the fields of both computational modeling and neuroimaging above and beyond the specific implications for the study of OL and arbitration. Ultimately, we show how a combination of pre-registered confirmatory analyses, supplemented by further (transparently reported) exploratory analyses, can facilitate substantial insight into the underlying mechanisms of observational learning and its arbitration.

**Figure 1.**
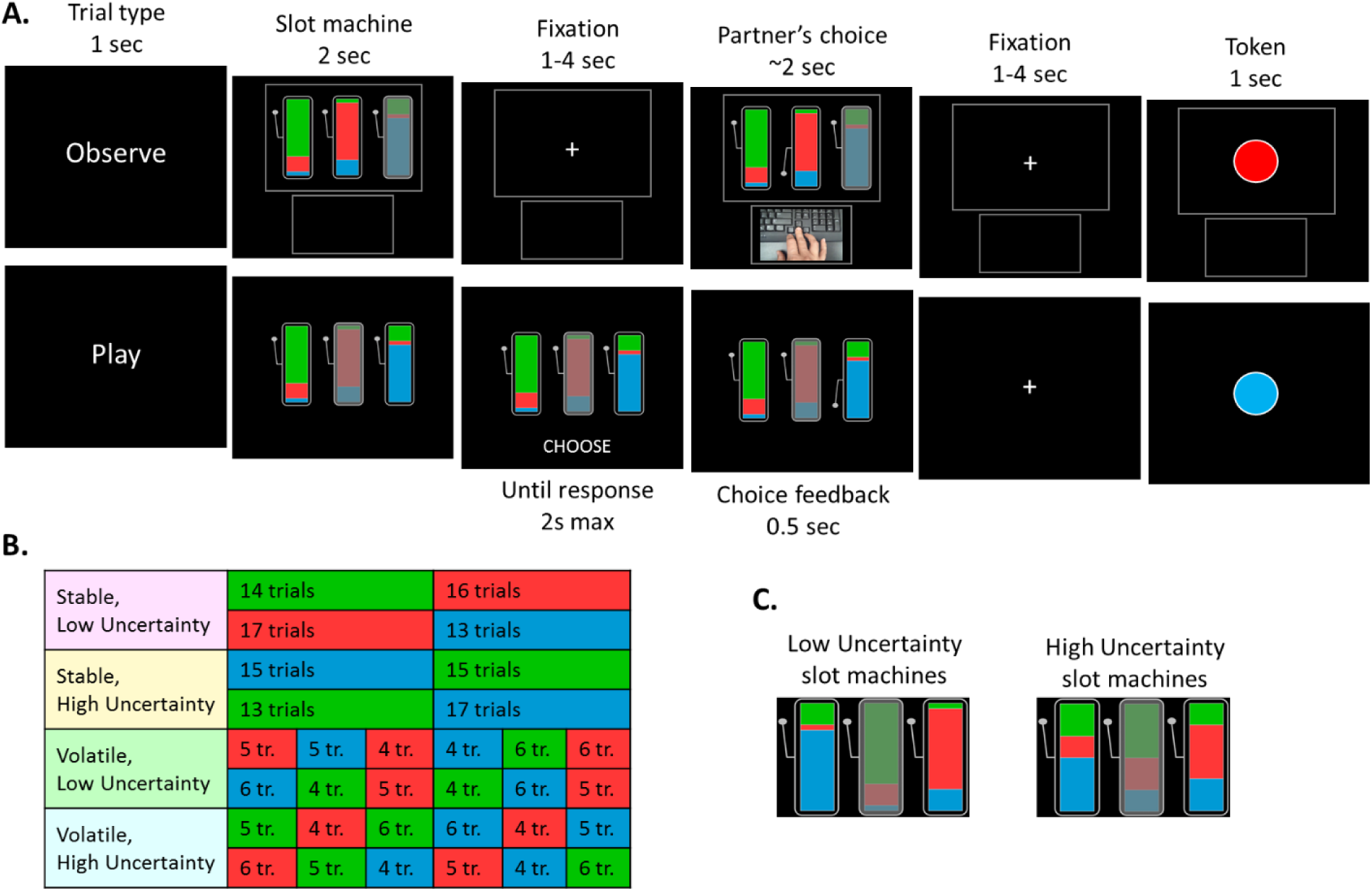
Observational Learning task design. (**A**) Illustration of two example trials, one where the participant observes another agent’s choice and one where the participant plays for themselves. In the observe trial the participant sees the slot machine choices available to the agent. The colors on each machine indicate the relative probability that a particular token will be delivered to the agent if that machine is chosen. The participant knows that the agent is provided with the information about which token is currently valuable. However, the participant is not directly informed about which token is currently valuable, instead they only get to observe the choice made by the agent. The agent’s choice is indicated by a video of the button press made by the agent and by the arm of the chosen slot machine being depressed. Crucially, the participant can infer which token is currently valuable through observing the agent’s choices (emulation), and computing the relative values of slot machines based on the observable color distributions. Alternatively, the participant can simply imitate the agent’s prior behavior on subsequent play trials, by choosing the action that was most frequently chosen by the agent in recent trials. (**B**) Outline of the task structure. The task contained 8 blocks of 30 trials, in a 2 (stable/volatile) by 2 (low/high uncertainty) design. The background color in the table depict which token is currently valuable (unknown to the participant). Block order was counterbalanced across subjects. In stable blocks, only one switch in valuable token occurred, roughly around the middle of the block. In volatile blocks, there were 5 switches. (**C**) Illustration of low vs high token uncertainty conditions. In low uncertainty blocks, the token probability distribution of the slot machines was [0.75, 0.2, 0.05], making value computation easier (less ambiguous) than for high uncertainty slot machines, for which the distribution was [0.5, 0.3, 0.2].

We predicted that participants’ observational learning behavior would be best explained by a mix of emulation and imitation, and that they would preferentially rely on one strategy over the other depending on the volatility and uncertainty in the environment. We also hypothesized that distinct neural signatures of imitation learning and emulation learning should be observed in the brain. Imitation was expected to recruit the fronto-parietal regions of the mirror neuron system involved in action observation, namely pre-motor cortex, inferior parietal cortex, and dlPFC ^8,10,12,31^. Because emulation involves inferring another agent’s goals and hidden mental state, it was therefore predicted to recruit regions of the mentalizing system ^16,32,33^. Finally, we hypothesized that the arbitration mechanism involved in tracking the relative reliability of the two strategies might depend on at least partially overlapping neural mechanisms as that used for arbitration in the experiential domain. Specifically, we hypothesized that the vlPFC and FPC would drive trial-by-trial variations in the arbitration controller ^27^, with the possible additional involvement of regions of the social brain, such as the TPJ.

## Results

In the task (**Fig. 1A**, see **Methods** for more details), participants learned to identify which of 3 tokens (green, blue or red) is valuable by observing another agent choose between explicit slot machines. The proportion of green, blue and red colors on each slot machine represents the probability of obtaining each token should that slot machine be chosen. Participants were instructed that the valuable token would switch many times throughout the task but weren’t told when the switches occurred. On 2/3 of trials (‘observe’ trials), they observed another agent play (through video) and knew that this other agent had full information about the valuable token and was therefore performing optimally. On 1/3 of trials (‘play’ trials), participants played for themselves. On each trial, one slot machine was unavailable – i.e. could not be chosen. Crucially, participants in this task can learn by inferring which token is currently valuable and computing the relative values of slot machines based on the observable color distributions (emulation). Alternatively, they can simply imitate the agent’s prior behavior by choosing the action that was most frequently performed by the agent in recent trials (imitation). By varying the position of the unavailable slot machine across trials, and particularly between observe trials and subsequent play trials, we were able to differentiate the two strategies behaviorally.

Importantly, we did not reveal the monetary value of the outcomes received by the observed agents to the participants. While the participants did see the color of the tokens obtained by the agents, they could not tell the current value of the token just from observing the token color obtained, ensuring that they had to utilize inference within their emulation system in order to work this out. Furthermore, because participants could not observe the actual reward amounts obtained by the agents, they could not utilize vicarious reward-learning, a third potential observational learning strategy in which one treats rewards obtained by another agent as if you had obtained them yourself and deploy a model-free reinforcement-learning strategy on those observed rewards to learn vicarious reward predictions ^4–6,34^. This critical feature of the task design ensured that we could study the effects of imitation (action learning) and emulation (goal inference) without the confounding effects of vicarious reinforcement-learning.

### Study 1

#### Behavioral signatures of imitation and emulation

A simple logistic regression was run to test for the presence of the two learning strategies. Specifically, choice of left versus right slot machine on each ‘play’ trial was predicted by an action learning regressor (signature of imitation: past left versus right actions performed by the partner) and a token learning regressor (signature of emulation: probability to choose left over right slot machine given inferred token information). Both regressors were found to significantly predict choice (action learning effect mean beta=0.865 ± 0.80 (SD), T_29_=5.94; token learning effect mean beta=1.174 ± 1.00 (SD), T_29_=6.42; all Ps<0.0001; **Fig. 2A**). This suggests that behavior on the task is a combination of imitation and emulation strategies.

**Figure 2.**
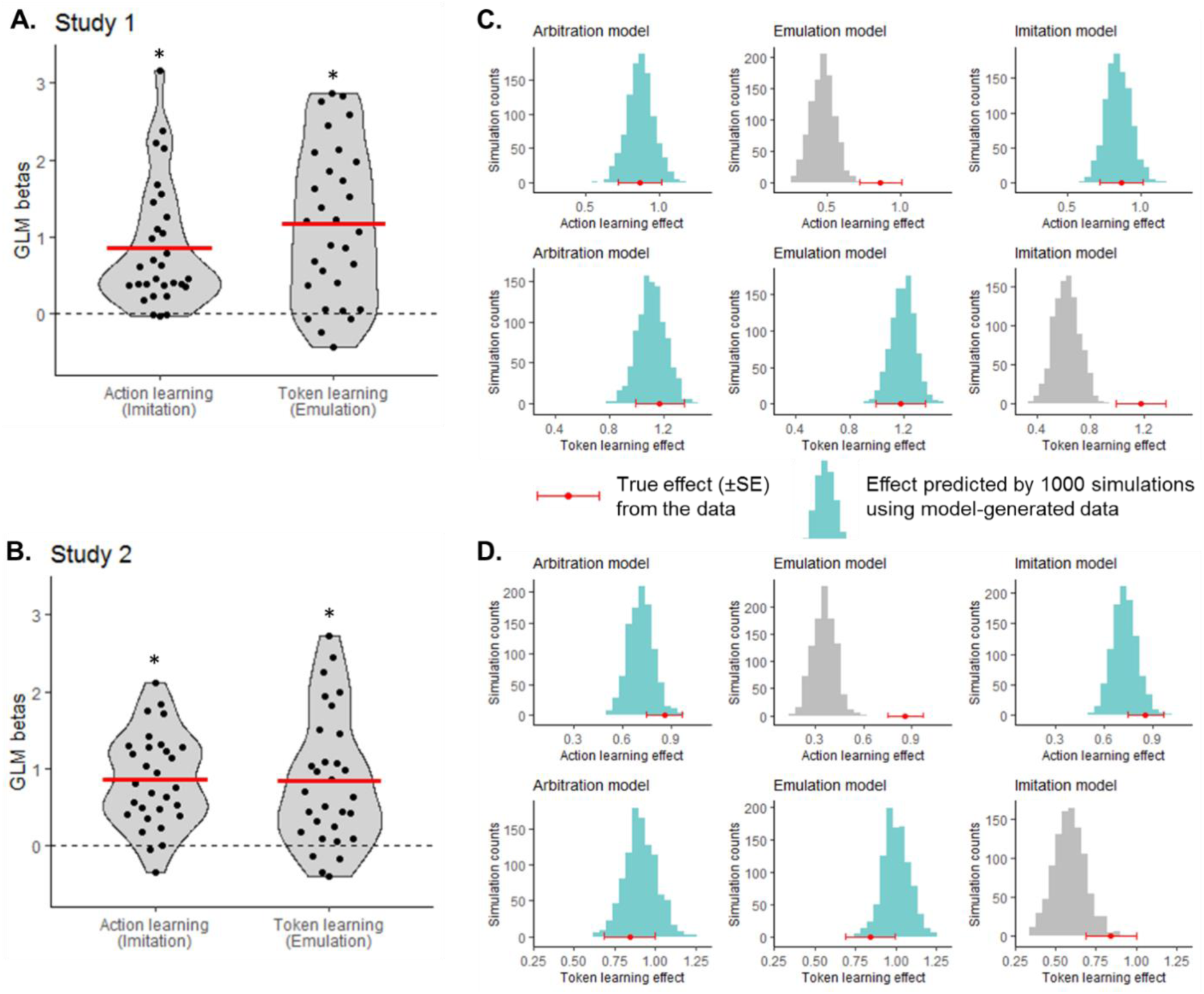
Behavioral signatures of imitation and emulation are captured by a computational model of arbitration. (**A-B**) A simple logistic regression was run to test for the presence of the two learning strategies. Choice was predicted by two regressors capturing the effect of past actions and the effect of past token inference. In both Study 1 (**A**) and Study 2 (**B**), both regressors had a significant effect on choice, suggesting a hybrid behavior between imitation and emulation learning. Dots represent individual subjects; the red bar represents the mean beta value. T-test: * P<0.0001; results were also confirmed using non-parametric permutation tests. (**C-D**) We then tested how well the winning computational model (Arbitration Model 7), as well as simple emulation (Model 2) and imitation (Model 3) models were able to capture the two behavioral effects obtained by the simple logistic regression presented in **A-B**: the action learning effect (top panels) and the token learning effect (bottom panels). The red data point above the X-axis depicts the true effect from the data, and the histogram shows the distribution of the recovered effects from the model-generated data. Effects that are well recovered are shown in light blue, while effects that are not well recovered are shown in gray. In both Study 1 (**C**) and Study 2 (**D**), the arbitration model (left panels) effectively captured both learning effects. In contrast, data generated by the emulation model (middle panels) only captured token-based learning, while data generated by the imitation model (right panels) only captured action-based learning.

#### Computational model of arbitration between imitation and emulation

To test whether this hybrid behavior between the two strategies can be explained by an arbitration mechanism, we performed computational modelling analyses. Specifically, we tested a set of 9 models, split into 5 classes (see **Methods** for details): emulation-only models (Models 1 and 2), which rely on multiplicative inference over token values; imitation-only models (Models 3 and 4), which use a reinforcement learning (RL) mechanism to learn about the other agent’s past actions; emulation RL models (Models 5 and 6), implemented as an RL mechanism rather than multiplicative inference; arbitration models (Models 7 and 8), in which the likelihood of relying on one strategy over the other varies as a function of each strategy’s relative reliability; and an outcome RL model (Model 9) to test the possibility that participants mistakenly learn from the token that is presented at the end of the trial. Two approaches were used to perform model comparison: between subjects out-of-sample predictive accuracy and group-level integrated Bayesian Information Criteria (iBIC ^35,36^). Arbitration models were found to perform best (**Table 1**). Specifically, Model 7 exhibited the highest out-of-sample accuracy (76.5%) and the lowest BIC. This suggests that an arbitration mechanism between imitation and emulation, based on the relative reliability of each strategy, explained behavior on the task better than each strategy individually or alternative models.

**Table 1.** Computational model comparison. Out-of-sample (OOS) accuracy was calculated in a 5-fold cross-validation analysis by estimating mean parameters in 24 subjects (4 groups of 6) and calculating the accuracy of predicted behavior on the remaining group of 6 subjects. Group-level integrated Bayesian Information Criteria (iBIC) were calculated following hierarchical model fitting. See **Methods** for details. Numbers in bold represent the winning model out of preregistered models (1-9). Numbers in bold and italics represent the winning model across all models, including exploratory models 10-11.

|  | Class | Model # | Parameters | OOS accuracy (%) |  | Group-level iBIC |  |
| --- | --- | --- | --- | --- | --- | --- | --- |
|  |  |  |  | Study 1 | Study 2 | Study 1 | Study 2 |
| Preregistered models | Emulation inference | 1 | $\beta$ | 67.8 | 66.1 | 2416 | 2458 |
| | | 2 | $\beta, \lambda$ | 68.3 | 67.9 | 2384 | 2368 |
| | Imitation RL | 3 | $\beta, \alpha$ | 71.3 | 71.6 | 2448 | 2455 |
| | | 4 | $\beta, \eta, \alpha_0$ | 70.5 | 69.8 | 2493 | 2472 |
| | Emulation RL | 5 | $\beta, \alpha$ | 67.1 | 67.4 | 2610 | 2537 |
| | | 6 | $\beta, \eta, \alpha_0$ | 65.4 | 64.8 | 2724 | 2597 |
| | Arbitration | 7 | $\beta_{em}, \beta_{im}, \delta, \alpha$ | **76.5** | **74.9** | **2296** | 2321 |
| | | 8 | $\beta_{em}, \beta_{im}, \delta, \alpha, \lambda$ | 73.9 | 72.1 | 2367 | **2291** |
| | Outcome RL | 9 | $\beta, \alpha$ | 58.7 | 58.5 | 3046 | 3052 |
| Exploratory model | Arbitration with 1-step IM | 10 | $\beta_{em}, \beta_{im}, \delta$ | **76.5** | **76.2** | **2236** | **2241** |

Finally, we tested whether the winning arbitration model could reliably recover the behavioral signatures of imitation and emulation identified above (i.e. the logistic regression effects shown in **Fig. 2A**). To do so, we generated behavioral data for each subject using the winning arbitration model (Model 7), as well as emulation Model 2 and imitation Model 3. Running the same logistic regression on the model-generated data (1000 iterations), we found that the arbitration model can reliably predict both action learning and token learning effects (**Fig. 2C left panel**). In contrast, the emulation model only predicts token learning (**Fig. 2C middle panel**) and the imitation model only predicts action learning (**Fig. 2C right panel**). These analyses show that the winning model is able to generate the behavioral effects of interest ^37^ and confirm the validity and specificity of our model.

#### Arbitration is influenced by uncertainty and volatility

Two factors were manipulated throughout the task (see **Methods** for details): volatility, consisting of high versus low frequency of switches in valuable token during a block (**Fig. 1B**) and uncertainty – the token probability distribution – associated with the slot machines (**Fig. 1C**). In volatile blocks, the actions performed by the partner becomes less consistent – we therefore refer to this manipulation as “action volatility” and predicted that volatility would predominantly tax the imitation system and indirectly favor emulation. Uncertainty in the token probability distribution makes inferring the best decision given the valuable token more difficult, while having no effect on the consistency of the partner’s actions – we therefore refer to this manipulation as “token uncertainty” and predicted that high uncertainty would tax the emulation system and indirectly favor imitation.

To test these predictions, we ran two analyses. First, we extracted the arbitration weight ω(t) values predicted using the best-fitting model parameters for each subject. These weight values, representing the probability of emulation (over imitation) for a given trial, were averaged for the two conditions of interest: volatile, low uncertainty trials, where we predict emulation should be maximal, and stable, high uncertainty trials, where we predict imitation should be maximal. As expected, the arbitration weight was higher in volatile, low uncertainty trials (mean ω=0.604 ± 0.26 (SD)) than on stable, high uncertainty trials (mean ω=0.474 ± 0.25 (SD); T_29_=15.22, P<0.0001; **Fig. 3A**). Across all 4 conditions (analyzed in a 2-by-2 repeated-measures ANOVA), there was also a main effect of volatility (F(1,29)=61.2, P<0.0001) and a main effect of uncertainty (F(1,29)=267.3, P<0.0001), suggesting a moderating effect of both manipulations.

**Figure 3.**
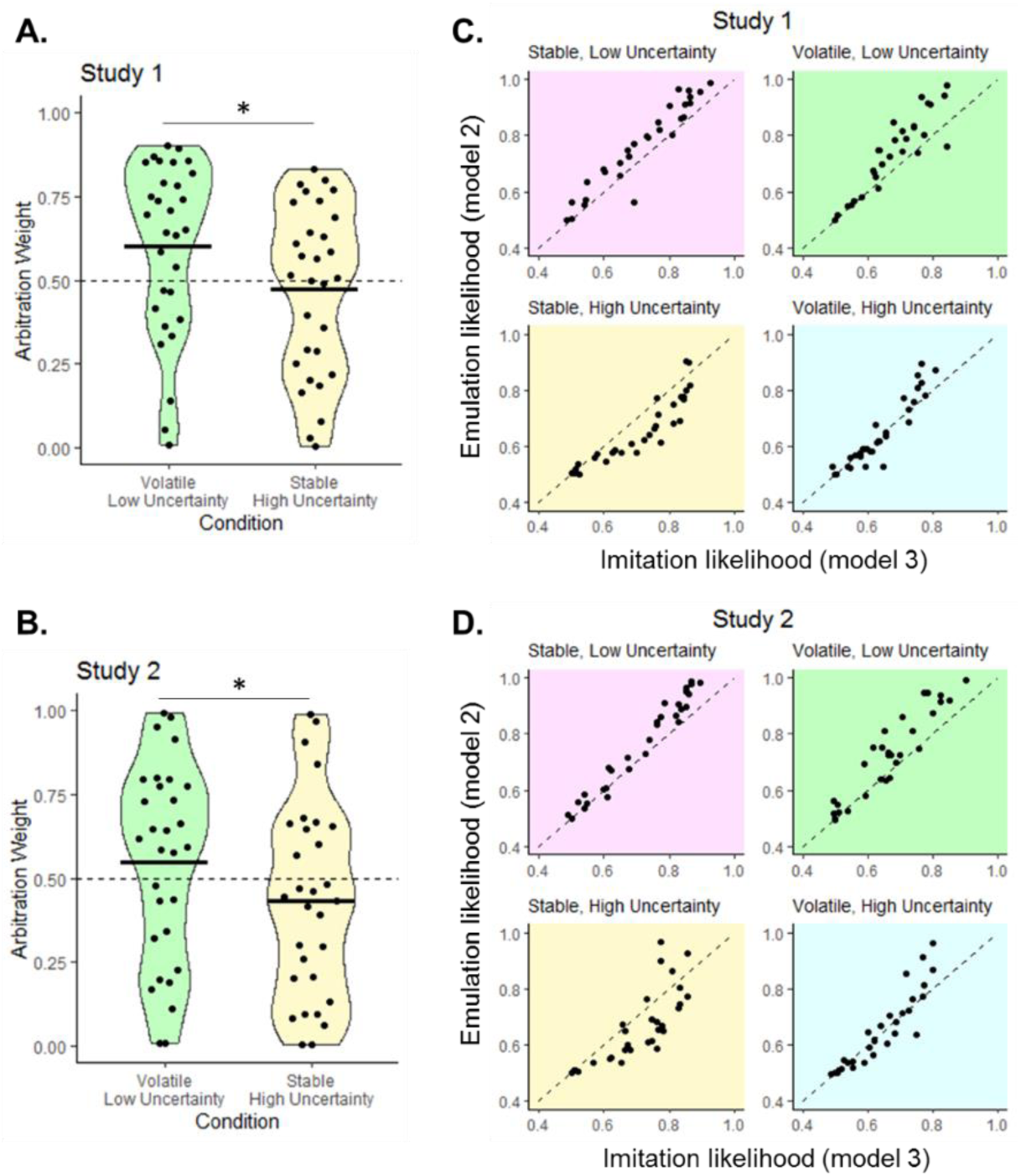
Modulation of arbitration by volatility and uncertainty. (**A-B**) Arbitration weight values, which represent the probability of relying on emulation over imitation on a given trial, were extracted from the winning arbitration model for each trial and averaged for each subject and each condition. The plots show, for Study 1 (**A**) and Study 2 (**B**), the mean and distribution of the arbitration weight for two conditions of interest: volatile/low uncertainty trials (green), and stable/high uncertainty trials (yellow). Dots represent individual subjects and the black bar represents the mean arbitration weight value for each condition. T-test: * P<0.0001; results were also confirmed using non-parametric permutation tests. (**C-D**) Mean per-trial emulation and imitation likelihood, extracted from Model 2 and Model 3 respectively, are plotted against each other separately for each of the 4 task conditions, showing that in both Study 1 (**C**) and Study 2 (**D**), most participants favor emulation (dots above the diagonal) when uncertainty is low (green & pink plots) but switch to imitation (dots below the diagonal) when the environment is stable and uncertainty is high (yellow plot).

Second, we compared the performance of the imitation-only model (Model 3) and the emulation-only model (Model 2), by calculating the mean likelihood per trial of each model and for each subject, and plotting them against each other separately for each condition (**Fig. 3C**). Dots above the diagonal represent participants who favor emulation, which is the case when token uncertainty is low (emulation-imitation likelihood difference for stable, low uncertainty condition = 0.051 ± 0.047 T_29_=5.88; for volatile, low uncertainty condition = 0.059 ± 0.061, T_29_=5.27; all Ps<0.0001), while dots below the diagonal mean that imitation is favored, which occurs when the partner’s actions are stable and token uncertainty is high (emulation-imitation likelihood difference for stable, high uncertainty condition = −0.053 ± 0.053, T_29_=−5.49, P<0.0001). There was no difference between imitation and emulation performance in volatile, high uncertainty trials (emulation-imitation difference = 0.007 ± 0.048, T_29_=0.74, P=0.46).

Taken together, these analyses suggest that emulation is preferred when action volatility is high (making action imitation learning more difficult) and when token uncertainty is low (making emulation value computation easier); while imitation is preferred in the opposite situation.

#### fMRI analyses of Study 1

Two fMRI models were utilized in the analysis of the of the Study 1 fMRI data: SPM GLM1 and SPM GLM2 (see **Methods** for details). Regressors were derived from each subject’s best fitting parameters from the winning arbitration Model 7, allowing to test for the presence of three types of signals. Emulation-related signals included trial-by-trial emulation reliability, update of token values (KL divergence) at the time of feedback, and entropy over token values at the time of initial slot machine presentation during observe trials. Imitation-related signals included trial-by-trial imitation reliability, and imitation action value difference at the time of initial slot machine presentation during observe trials. Finally, arbitration-related signals included the trial-by-trial difference in reliability (emulation – imitation), which is assumed to drive arbitration, and the chosen action value at the time of choice on play trials. The effect of most of these regressors were assessed in SPM GLM1. SPM GLM2 tested for the separate effects of imitation reliability and emulation reliability instead of the reliability difference regressor.

Eight regions of interests (ROIs) were defined based on previous literature on observational learning, social inference and arbitration processes during learning ^15,27^ (see **Methods** for details). The effect of the different regressors were assessed in each ROI by extracting the mean signal across all voxels in the ROI for each subject, then averaging across subjects (**Table S1**). T-tests were performed to establish significance, and were confirmed with non-parametric permutation tests (with 10,000 permutations) in all ROI analyses, since data were not always normally distributed across the samples. Because the goal of this ROI analysis was to generate hypotheses to be confirmed in Study 2, we did not correct for multiple comparisons across the different ROIs. Instead, in our subsequent pre-registration for Study 2, we selected significant ROIs in Study 1 to restrict the space of regions to examine in Study 2. In addition to the ROI analysis, whole-brain group-level T-maps were evaluated, and uploaded on NeuroVault (https://neurovault.org/collections/UBXVWSMN/), before the collection of Study 2 data. Significant activation clusters for Study 1 are reported in **Table S2**. Significant clusters were identified and saved as functional regions of interest for later examination in Study 2. Results derived from both analyses are presented in **Fig. 4** for arbitration signals, **Fig. 5** for emulation signals, and **Fig. 6** for imitation signals.

**Figure 4.**
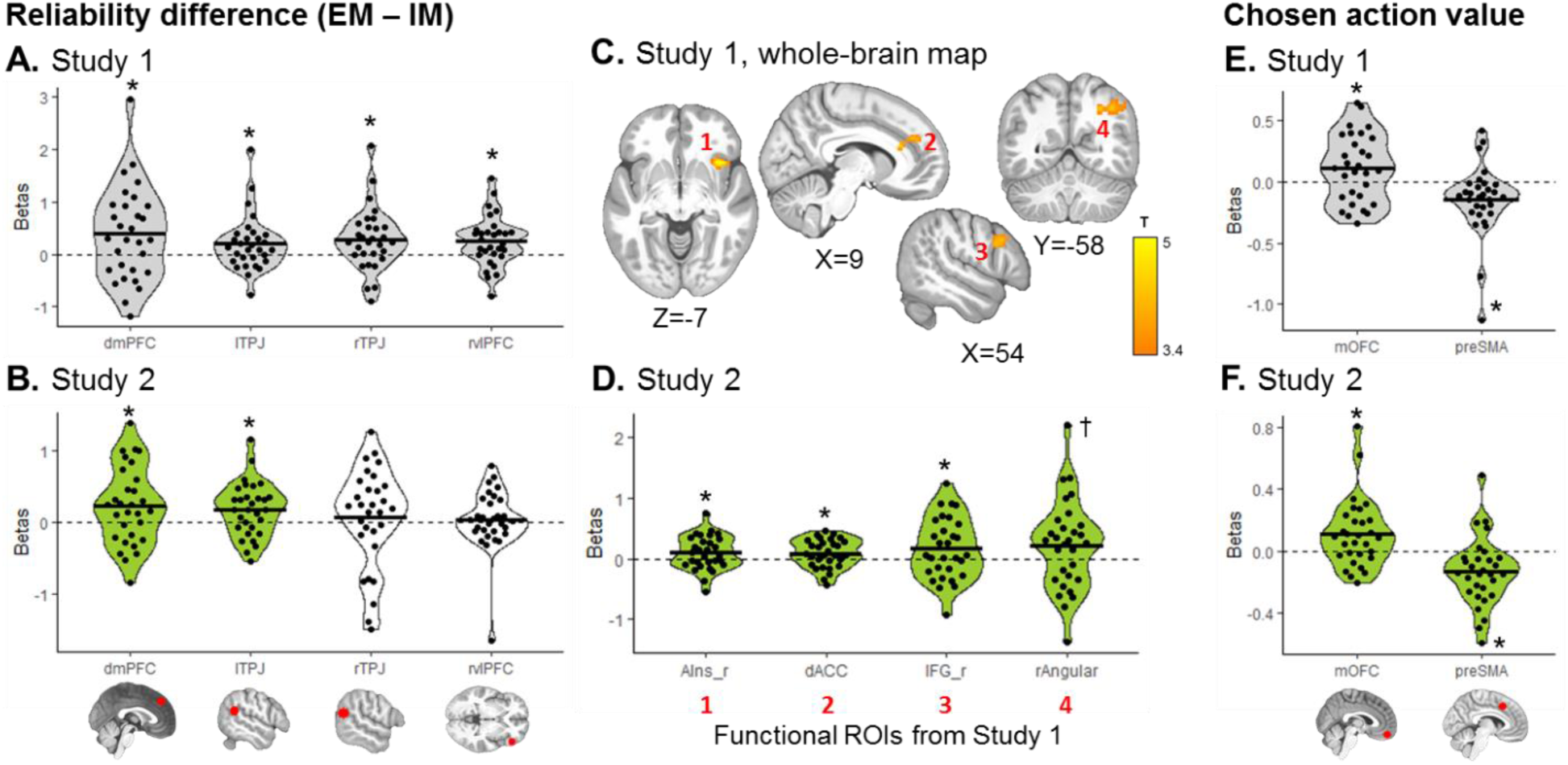
Arbitration signals, pre-registered analyses. For each arbitration contrast of interest in SPM GLM1 – reliability difference (**A-D**) and chosen action value (**E-F**) – mean signal was extracted from each pre-registered ROI. (**A, E**) Regions with significant signals in Study 1, plotted in grey, were selected as hypotheses and a priori ROIs for Study 2. (**C**) Whole-brain maps for the reliability difference signal were also examined in Study 1, with a cluster-forming threshold of P<0.001 uncorrected, followed by cluster-level FWE correction at P<0.05. Significant clusters were then saved as functional ROIs to be examined in Study 2. Note that there was no cluster surviving correction for the chosen value signal. (**B, D, F**) Green plots represent significant effects in Study 2, confirming the a priori hypothesis from Study 1. White plots represent hypotheses that were not confirmed in Study 2. Dots represent individual subjects and the black bar represents the mean beta estimate for each effect. T-tests: * P<0.05, ^†^ P=0.052. The same results were found using non-parametric permutation tests.

**Figure 5.**
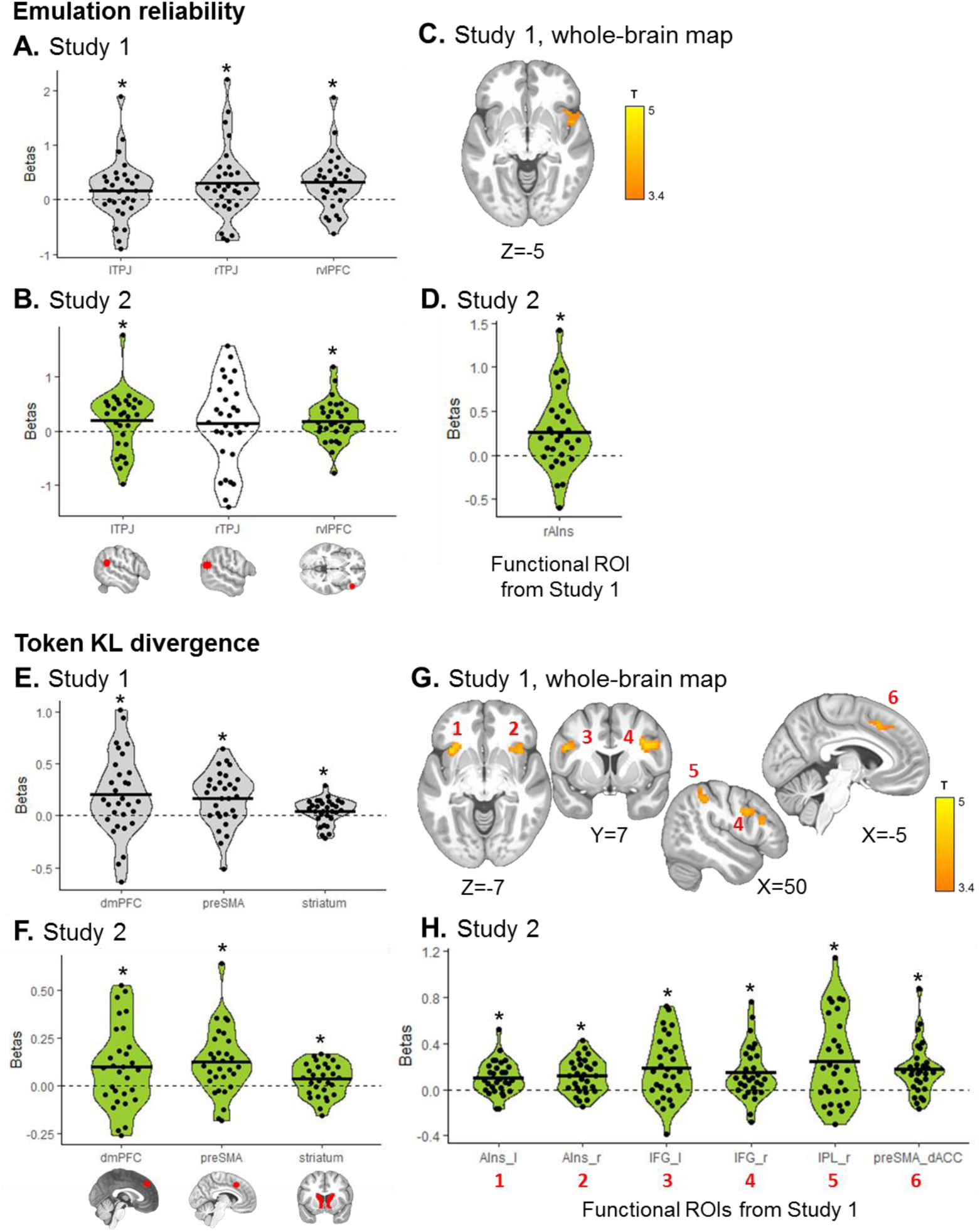
Emulation signals, pre-registered analyses. For each emulation contrast of interest – emulation reliability in SPM GLM2 (**A-D**) and update in token values in SPM GLM1 (**E-H**) – mean signal was extracted from each pre-registered ROI. (**A, E**) Regions with significant signals in Study 1, plotted in grey, were selected as hypotheses and a priori ROIs for Study 2. (**C, G**) Whole-brain maps for each contrast were also examined in Study 1, with a cluster-forming threshold of P<0.001 uncorrected, followed by cluster-level FWE correction at P<0.05. Significant clusters were then saved as functional ROIs to be examined in Study 2. (**B, D, F, H**) Green plots represent significant effects in Study 2, confirming the a priori hypothesis from Study 1. White plots represent hypotheses that were not confirmed in Study 2. Dots represent individual subjects and the black bar represents the mean beta estimate for each effect. T-tests: * P<0.05. The same results were found using non-parametric permutation tests.

**Figure 6.**
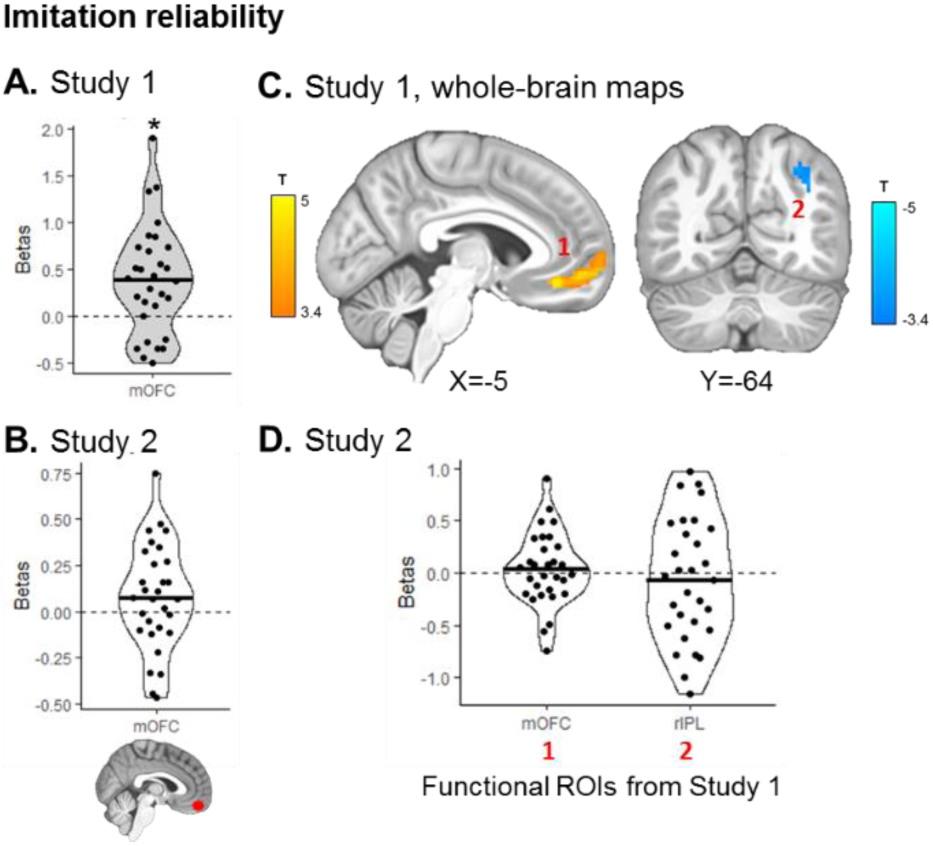
Imitation signals, pre-registered analyses. Mean imitation reliability signal was extracted from each pre-registered ROI in SPM GLM2. (**A**) The medial OFC ROI was found to significantly track imitation reliability in Study 1 (* P<0.05, t-test and non-parametric permutation test), and was selected as an a priori ROI for Study 2. (**C**) Whole-brain maps for positive (orange scale) and negative tracking (blue scale) of imitation reliability were also examined in Study 1, with a cluster-forming threshold of P<0.001 uncorrected, followed by cluster-level FWE correction at P<0.05. Significant clusters were then saved as functional ROIs to be examined in Study 2. (**B, D**) The a priori hypotheses were not confirmed in Study 2.

The difference in reliability between imitation and emulation, predictive of trial-by-trial arbitration tendencies, was reflected in the activity of four ROIs (**Fig. 4A**): dmPFC (mean beta = 0.383 ± 0.89 (SD), T_29_= 2.37, P=0.012), bilateral TPJ (left: mean beta = 0.201 ± 0.53 (SD), T_29_=2.10, P=0.022; right: mean beta = 0.277 ± 0.59 (SD), T_29_=2.56, P=0.008) and right vlPFC (mean beta = 0.250 ± 0.48 (SD), T_29_=2.86, P=0.004). Using whole-brain group analyses, four significant clusters were found (all P_FWE_<0.05; **Fig. 4C**): right anterior insula (peak voxel: 40, 17, −12; T_29_=5.97), dorsal ACC, partially overlapping with the dmPFC ROI (peak voxel: 13, 44, 26; T_29_=4.80), right IFG (peak voxel: 45, 4, 21; T_29_=5.02), and right angular gyrus (peak voxel: 40, −74, 48; T_29_=4.38). At the time of choice, the expected value of the chosen slot machine was coded positively in the mOFC (mean beta = 0.110 ± 0.28 (SD), T_29_=2.16, P=0.019) and negatively in the pre-supplementary motor area (preSMA; mean beta = −0.144 ± 0.29 (SD), T_29_=−2.74, P=0.005; **Fig. 4E**). There was no cluster surviving correction for chosen action value in the whole-brain analysis.

Emulation reliability was represented in the bilateral TPJ (left: mean beta = 0.172 ± 0.55 (SD), T_29_=1.72, P=0.048; right: mean beta = 0.299 ± 0.66 (SD), T_29_=2.46, P=0.010) and right vlPFC (mean beta = 0.320 ± 0.50 (SD), T_29_=3.50, P=0.0008; **Fig. 5A**). In the whole-brain analysis, an additional cluster was identified encoding emulation reliability in the right anterior insula (peak voxel: 43, 17, −12; T_29_=4.82, P_FWE_<0.05; **Fig. 5C**). Update in token values, a key signature of emulation learning calculated as the KL divergence between prior and posterior token values, was tracked at the time of feedback (during observation of the partner’s action) in three regions (**Fig. 5E**): dmPFC (mean beta = 0.201 ± 0.39 (SD), T_29_=2.84, P=0.004), preSMA (mean beta = 0.170 ± 0.26 (SD), T_29_=3.62, P=0.0006) and dorsal striatum (mean beta = 0.043 ± 0.12 (SD), T_29_=1.99, P=0.028). The whole brain analysis revealed significant clusters (all P_FWE_<0.05; **Fig. 5G**) tracking emulation update in the bilateral anterior insula (left: peak voxel −33, 14, −10; T_29_=4.88; right: peak voxel 40, 19, −2; T_29_=4.65), bilateral IFG (left: peak voxel −48, 7, 26; T_29_=4.41; right: peak voxel 35, 9, 33; T_29_=4.69), right supramarginal and inferior parietal cortex (peak voxel: 53, −39, 46; T_29_=4.09) and preSMA extending into the dorsal ACC (peak voxel: −8, 19, 46; T_29_=4.44).

Finally, imitation reliability was found to be significantly tracked in the medial OFC ROI (mean beta = 0.387 ± 0.58 (SD), T_29_=3.67, P=0.0005, **Fig. 6A**) and in a significant cluster spanning over the medial OFC and vmPFC (peak voxel: 3, 37, −7; T_29_=5.01, P_FWE_<0.05; **Fig. 6C**) Imitation reliability was also negatively tracked in a right inferior parietal cluster (peak voxel: 48, −46, 58; T_29_=4.43, P_FWE_<0.05; **Fig. 6C**).

For completeness, all the pre-registered ROI results are reported in **Table S1** and **Figure S1**, and whole-brain analyses in **Table S2**. The statistical significance of all ROI results remained unchanged when using non-parametric permutation tests.

#### Pre-registration for Study 2

In order to evaluate the extent to which our computational model fitting and fMRI results could be replicated in an independent sample, which is the gold standard method to establish the extent to which one’s findings are not susceptible to overfitting ^38–41^, we pre-registered our computational models as designed and utilized for Study 1, our computational model-fitting procedures to behavior, our fMRI analysis pre-processing pipeline, and our fMRI statistical models and utilized this exact pipeline for the analysis of Study 2 (project: https://osf.io/49ws3/; pre-registration: https://osf.io/37xyq). For the case of Study 2 fMRI results, we used the 8 ROIs described earlier, as well as a subset of the clusters identified in Study 1 that were saved as functional regions of interest for use in Study 2. We focused specifically on testing the replicability of the findings reported in Study 1 in both the behavioral and neuroimaging data.

### Study 2

#### Replication of behavioral and computational modelling results

The simple logistic regression testing for the presence of both token learning and action learning strategies yielded the same findings. As in Study 1, both regressors were also found to significantly predict choice in Study 2 (action learning effect: mean beta=0.857 ± 0.60 (SD), T_29_=7.78; token learning effect: mean beta=0.843 ± 0.85 (SD), T_29_=5.42; both Ps<0.0001; **Fig. 2B**).

The computational modelling results revealed that the arbitration Model 7 also had the highest out-of-sample accuracy of all pre-registered models (74.9%; **Table 1**). While Model 7 had the lowest iBIC in Study 1, Model 8 had the lowest iBIC in Study 2 (**Table 1**). Models 7 and 8 are both arbitration models and thus very similar. The only difference between the two models is the presence of a free parameter λ (in Model 8), which represents trust in current token values and captures the tendency of participants to overestimate volatility in the environment (see Methods for details). Given that Model 7 is a more parsimonious model, we kept it as our winning model in both Study 1 and Study 2, thus maintaining consistency across studies. We also confirm that data generated using arbitration Model 7 in Study 2 can, similarly to Study 1, reliably predict both action learning and token learning effects (**Fig. 2D**). As a sanity check, we find a very similar pattern of results when using arbitration Model 8 to generate data (**Fig. S2**).

Finally, arbitration was influenced by uncertainty and volatility in the same way in Study 2 as in Study 1. Action volatility was found to increase the arbitration weight (extracted from Model 7), while token uncertainty decreased it (high volatility & low uncertainty: mean ω=0.550 ± 0.29 (SD); low volatility & high uncertainty: mean ω=0.434 ± 0.29 (SD); difference: T_29_=10.97, P<0.0001; **Fig. 3B**). Across all 4 conditions, there was also a main effect of volatility (F(1,29)=47.3, P<0.0001) and a main effect of uncertainty (F(1,29)=124.8, P<0.0001), confirming the combined effect of both manipulations. Comparing the performance of the imitation-only and emulation-only models (**Fig. 3D**), we also replicated the findings from Study 1. Emulation was favored when token uncertainty is low, as shown by significantly positive emulation-imitation likelihood difference (stable, low uncertainty condition = 0.050 ± 0.043 (SD), T_29_=6.31; volatile, low uncertainty condition = 0.061 ± 0.061 (SD), T_29_=5.49; all Ps<0.0001). Imitation was favored when token uncertainty is high and partner’s actions are stable, as shown by significantly negative emulation-imitation likelihood difference in that condition (mean = −0.045 ± 0.081 (SD), T_29_=−3.04, P=0.0025). There was also no difference between imitation and emulation in volatile, high uncertainty trials (mean = 0.009 ± 0.059 (SD), T_29_=0.90, P=0.37).

Overall, these findings show in both studies (i) that participants combine two learning strategies to perform the task, (ii) that this hybrid behavior is best explained by an arbitration model between imitation and emulation, and (iii) that participants flexibly adapt their learning strategy depending on the environment.

#### Replication of emulation and decision value signals, but not imitation signals

BOLD responses related to the emulation strategy were largely replicated in Study 2. Specifically, emulation reliability was found to be significantly represented in two of the three ROIs identified in Study 1 (**Fig. 5B**) – the left TPJ (mean beta = 0.195 ± 0.55 (SD), T_29_=1.96, P=0.030) and the right vlPFC (mean beta = 0.186 ± 0.39 (SD), T_29_=2.62, P=0.0069), but not in the right TPJ (mean beta = 0.137 ± 0.78 (SD), T_29_=0.96, P=0.17). Emulation reliability was also significant in the right anterior insula functional ROI saved from Study 1’s whole-brain map (mean beta = 0.258 ± 0.43 (SD), T_29_=3.28, P=0.0014; **Fig. 5D**). The KL divergence over token values was tracked in the same three ROIs (**Fig. 5F**), namely dmPFC (mean beta = 0.098 ± 0.21 (SD), T_29_= 2.52, P=0.0087), preSMA (mean beta = 0.123 ± 0.17 (SD), T_29_=3.91, P=0.00025) and dorsal striatum (mean beta = 0.033 ± 0.085 (SD), T_29_=2.15, P=0.020). Examining functional clusters saved from Study 1, all six regions also showed significant emulation update signal in Study 2 (**Fig. 5H**): bilateral anterior insula (left: mean beta = 0.102 ± 0.15 (SD), T_29_=3.80, P=0.0003; right: mean beta = 0.117 ± 0.15 (SD), T_29_=4.41, P<0.0001), bilateral IFG (left: mean beta = 0.187 ± 0.27 (SD), T_29_=3.74, P=0.0004; right: mean beta = 0.147 ± 0.23 (SD), T_29_=3.49, P=0.0008), right inferior parietal extending into supramarginal cortex (mean beta = 0.246 ± 0.38 (SD), T_29_=3.56, P=0.0007), and preSMA/dorsal ACC (mean beta = 0.174 ± 0.22 (SD), T_29_=4.30, P<0.0001). Entropy over token values, at the time of initial slot machine presentation on observe trials, was also found to be negatively represented in the mOFC (Study 1: mean beta = −0.080 ± 0.25 (SD), T_29_=−1.73, P=0.047; Study 2: mean beta = −0.107 ± 0.19 (SD), T_29_=−3.08, P=0.0023; **Fig. S1A-B**), suggesting the mOFC is more active when token values are more certain.

Decision values signals, calculated as the expected value of the chosen slot machine on play trials, recruited the same ROIs in Study 2 (**Fig. 4F**), with positive value coding in mOFC (mean beta = 0.109 ± 0.22 (SD), T_29_=2.70, P=0.0057) and negative value coding in preSMA (mean beta = −0.135 ± 0.22 (SD), T_29_=−3.38, P=0.0011). The reliability difference between the two strategies, assumed to drive the arbitration process, was found to replicate in two of the four ROIs identified in Study 1 (**Fig. 4B**) – the left TPJ (mean beta = 0.182 ± 0.38 (SD), T_29_=2.63, P=0.0068) and the dmPFC (mean beta = 0.228 ± 0.54 (SD), T_29_=2.31, P=0.014) – as well as in functional clusters in the dorsal ACC (mean beta = 0.089 ± 0.23 (SD), T_29_=2.09, P=0.023), right anterior insula (mean beta = 0.099 ± 0.28 (SD), T_29_=1.95, P=0.031), IFG (mean beta = 0.173 ± 0.51 (SD), T_29_=1.87, P=0.036) and angular gyrus, at trend level (mean beta = 0.225 ± 0.74 (SD), T_29_=1.67, P=0.052; **Fig. 4D**).

However, when examining more closely whether this signal was a true difference signal, by separately extracting emulation and imitation reliabilities from these ROIs, we did not find robust evidence for negative tracking of imitation reliability or significant representation of the two reliability signals in opposite directions in either Study 1 or 2 (**Fig. S3**). Instead, reliability difference signals were mainly driven by positive tracking of emulation reliability, suggesting that the arbitration mechanism might rely more on emulation reliability than on imitation reliability, at least in so far as it is implemented in the brain. In addition, all signals pertaining to the imitation strategy, namely imitation reliability (all T_29_<1.49, all Ps>0.15; **Fig. 6B** and **6D**) and the difference in imitation action values (see **Fig. S1D** and **Table S2** for details) did not replicate well in Study 2. Given our experimental evidence, we concluded that the component of our original model focusing on imitation may not be as justified as the emulation component. Furthermore, the findings also suggested to us that the arbitration mechanism may rely less on imitation reliability than originally hypothesized. Armed with these conclusions, we decided to revisit our computational model and to re-analyze both behavioral and neuroimaging datasets in order to test the possibility of an alternative computational modeling strategy for imitation and for the arbitration between the two mechanisms. This new analysis is described as exploratory since it is distinct from the confirmatory analyses reported in Study 2. Importantly, however, because we are still able to test our new model and analysis on two independent datasets, we can confirm the robustness and replicability of our findings.

### Exploratory analyses: arbitration between emulation and simpler (1-step) imitation strategy

#### Behavioral evidence

Given that imitation-related signals were not reliably found across the two studies, we hypothesized that imitation learning may be implemented differently, both behaviorally and in the brain. Specifically, in our pre-registered analyses, we initially modelled action imitation as an RL model, which computes action values from multiple trials of experience, but imitation could be as simple as just copying the most recent action, rather than slowly computing values over time. We thus tested this possibility of a simpler imitation strategy (“1-step imitation”), whereby out of the two available actions on a given play trial, the action that was most recently performed by the partner is repeated (see **Methods** for details). Emulation action values and emulation reliability were computed as before, and arbitration was assumed to be driven by the reliability of emulation only. Thus, the resulting arbitration process was such that if the reliability of emulation is high, participants will be more likely to rely on emulation, whereas if it is low, they will be more likely to default to imitation.

In both studies, this new arbitration model (Model 10) was found to perform better than the pre-registered winning model (Model 7), with higher out-of-sample accuracy (Study 1: 76.5%, Study 2: 76.2%) and lower iBIC values (**Table 1**). Given that this simpler imitation strategy does not require a learning rate parameter, Model 10 is more parsimonious than Model 7, which could in part account for the lower iBIC values. However, the improved out-of-sample performance for Model 10 is a key result here, as it suggests that Model 10 affords better out-of-sample generalization than Model 7, indicating that the difference in model performance is not just due to model complexity alone. A possible explanation for this finding is that Model 7 was overfitting, or that Model 10 offers a better computational distinction between the two strategies. Using data generated by this new arbitration model, we were also able to recover both action learning and token learning effects obtained from a simple logistic regression analysis (**Fig S4**), thus confirming the validity of this new, more parsimonious model. This suggests that participants’ hybrid behavior on the task is better explained by an arbitration process between inferring the valuable token (emulation) and repeating the partner’s most recent action (1-step imitation) than by an arbitration process in which imitation is implemented as a reinforcement learning mechanism over the recent history of actions.

#### Neuroimaging evidence

The next question is whether neuroimaging evidence would support this proposed arbitration process. Specifically, this new arbitration model makes the following predictions. Trial-by-trial emulation reliability should be represented in the brain, given that it is assumed to drive the arbitration process and the likelihood of relying on emulation or defaulting to imitation. Learning signals specific to each strategy should be observed at the time of feedback, when the partner’s action is shown. For emulation, this update signal takes the form of the KL divergence between prior and posterior token value, as defined in the pre-registered analyses. For imitation, this signal was defined as tracking whether or not the partner’s current action repeats the most recent past action. Our prediction is that an update signal should occur when there is an action change, i.e. the most recent action is available on the current trial but the partner chooses a different option. To test these predictions, we defined an additional model of the fMRI BOLD signal, SPM GLM3 (see **Methods** for details).

Using Bayesian model selection (BMS, see **Methods** for details), we first confirmed that this new SPM model (SPM GLM3) performed better than the pre-registered model testing for neural signatures of imitation implemented as an RL mechanism (SPM GLM2). In both studies, GLM3 was associated with the highest exceedance probability averaged across all grey matter voxels (Study 1: 0.861; Study 2: 0.946) and with a vast majority of grey matter voxels with an exceedance probability higher than 0.75 (**Fig. S5**). When examining the average exceedance probabilities in our set of pre-registered ROIs, evidence was also overwhelmingly in favor of SPM GLM3 (**Table S3**). This indicates that this new SPM GLM, based on the best performing model of behavior, also explained trial-by-trial variations in BOLD signal best.

Arbitration in this new model is driven by variations in the reliability of emulation. We thus tested whether these variations were represented in the brain. Unsurprisingly, given that the calculation of emulation reliability did not change from the pre-registered analyses, we found very similar results, and robust representation of this signal driving arbitration between the two strategies. In both studies emulation reliability was found to be represented in the same three ROIs: the right vlPFC (Study 1: mean beta = 0.253 ± 0.48 (SD), T_29_=2.90, P=0.0035; Study 2: mean beta = 0.232 ± 0.45 (SD), T_29_=2.84, P=0.0041), the left TPJ (Study 1: mean beta = 0.154 ± 0.47 (SD), T_29_=1.80, P=0.041; Study 2: mean beta = 0.252 ± 0.62 (SD), T_29_=2.23, P=0.017), and the right TPJ, albeit only at trend level in Study 2 (Study 1: mean beta = 0.284 ± 0.62 (SD), T_29_=2.52, P=0.0088; Study 2: mean beta = 0.234 ± 0.78 (SD), T_29_=1.65, P=0.055; **Fig. 7A-B**). Exploratory conjunction analysis additionally revealed significant clusters in the ACC, bilateral insula, and supramarginal gyrus (**Fig. 7C** and **Table S4**).

**Figure 7.**
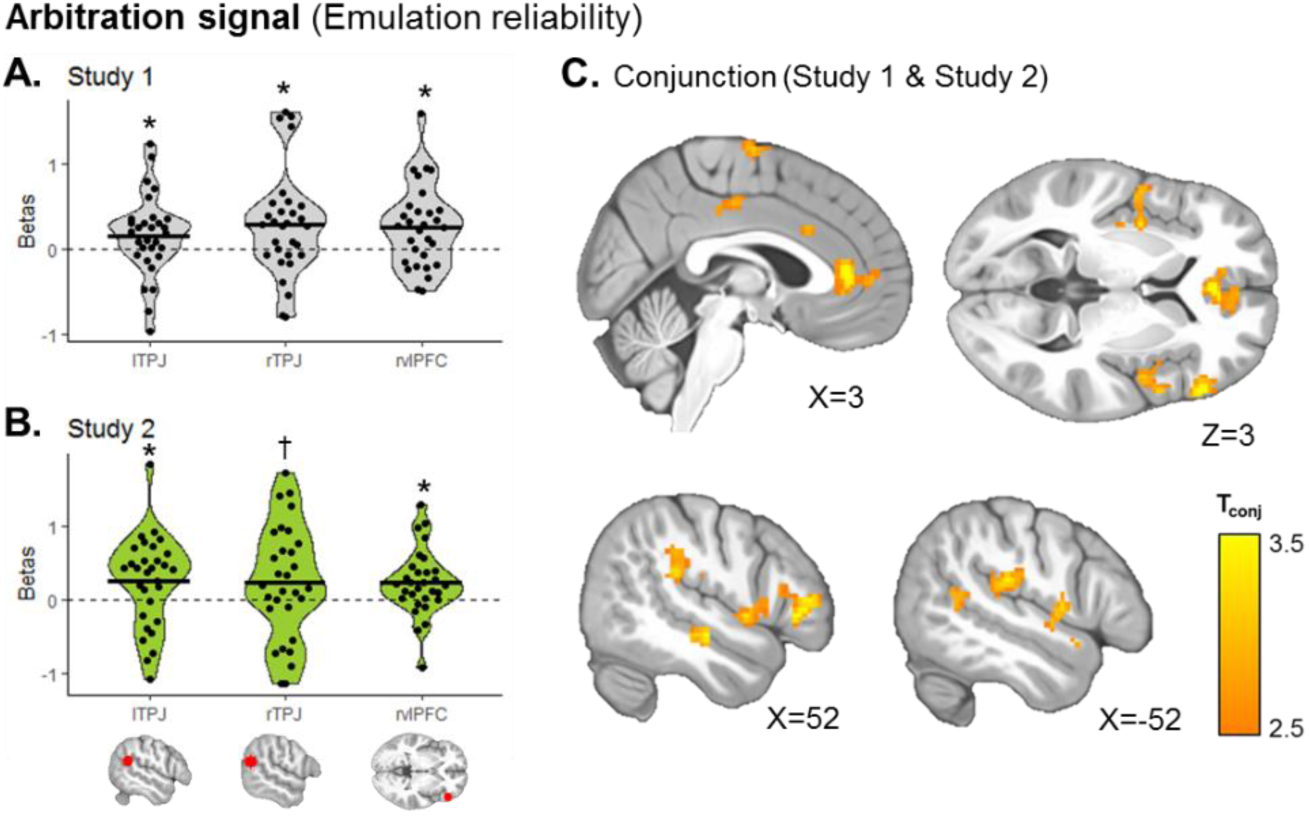
Neural representation of emulation reliability as an arbitration signal. Trial-by-trial values of the reliability of emulation, as predicted by the winning arbitration Model 10, were added as a parametric modulator of the BOLD signal during both observe and play trials. (**A-B**) Using our preregistered ROIs, we found in both Study 1 (**A**) and Study 2 (**B**) that this signal was represented in the bilateral TPJ and in the right vlPFC. Dots represent individual subjects; the black bar represents the mean beta value for each regressor. T-tests: * P<0.05, ^†^ P=0.055. The same results were found using non-parametric permutation tests. (**C**) Using exploratory whole-brain conjunction analysis between the second-level T-maps of Study 1 and Study 2, we show additional clusters tracking emulation reliability, including in the ACC and bilateral insula (see **Table S4** for details). Maps were thresholded at P_conjunction_<0.0001 uncorrected, followed by whole-brain cluster-level family-wise error correction at P<0.05 (cluster size ≥ 30).

We then examined update signals specific to each strategy at the time of feedback, when the participant observes the partner’s action and can update their estimates of token values and/or of which action is best to perform. As expected given the pre-registered analyses, the neural signature of emulation inference, calculated as the KL divergence over inferred token values, showed a very similar pattern to the pre-registered results, with significant effects in the dmPFC (Study 1: mean beta = 0.164 ± 0.34 (SD), T_29_=2.63, P=0.0067; Study 2: mean beta = 0.112 ± 0.23 (SD), T_29_=2.72, P=0.0054), preSMA (Study 1: mean beta = 0.135 ± 0.24 (SD), T_29_=3.09, P=0.0022; Study 2: mean beta = 0.093 ± 0.17 (SD), T_29_=2.96, P=0.0031), right TPJ, albeit only at trend in Study 2 (Study 1: mean beta = 0.062 ± 0.18 (SD), T_29_=1.84, P=0.038; Study 2: mean beta = 0.073 ± 0.24 (SD), T_29_=1.66, P=0.054), and dorsal striatum (Study 1: mean beta = 0.043 ± 0.12 (SD), T_29_=2.00, P=0.027; Study 2: mean beta = 0.028 ± 0.075 (SD), T_29_=2.07, P=0.024; **Fig. 8A-B**). Exploratory conjunction analysis confirmed these clusters, as well as additional clusters in the bilateral insula, inferior frontal gyrus, and other frontoparietal regions (**Fig. 8C**; see **Table S4** for details).

**Figure 8.**
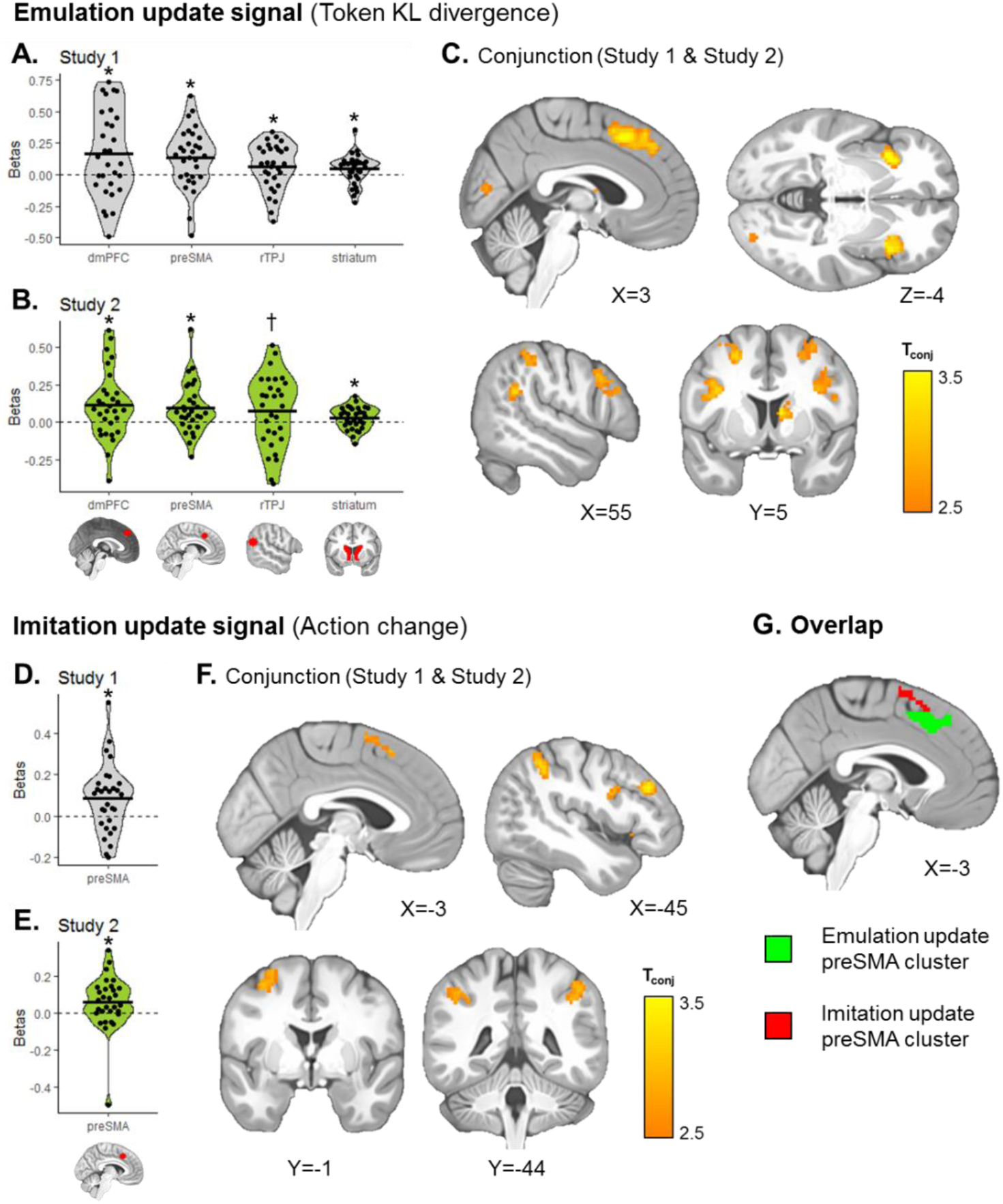
Imitation and emulation update signals during observation. KL divergence over token values (necessary for emulation learning) and changes in the partner’s action relative to the previous trial (necessary for imitation learning) were added as parametric modulators of the BOLD signal during feedback, when the partner’s action is shown. The two regressors competed for variance within the same model. (**A, B, D, E**) Using our preregistered ROIs, we found in both studies significant emulation update signals in the dmPFC, preSMA, right TPJ and dorsal striatum (**A-B**) and significant action imitation signals in the preSMA ROI (**D-E**). Dots represent individual subjects; the black bar represents the mean beta value for each regressor. T-tests: * P<0.05, ^†^ P=0.054. The same results were found using non-parametric permutation tests. (**C, F**) Using exploratory whole-brain conjunction analysis between the second-level T-maps of Study 1 and Study 2, we show additional clusters tracking emulation (**C**) or imitation (**F**) update (see **Table S4** for details). Maps were thresholded at P_conjunction_<0.0001 uncorrected, followed by whole-brain cluster-level family-wise error correction at P<0.05. (**G**) Given that both imitation and emulation update signals recruited the preSMA ROI, we tested for a possible overlap between the two signals and find that the emulation update signal (green) is more anterior and ventral than the imitation update signal (red) with no overlap between the two.

However, contrary to the pre-registered findings in which imitation signals were not replicated, here we find robust tracking of whether the partner’s current action marks a change or a repeat of the previous action, consistent with the simpler 1-step imitation strategy. This action change imitation signal was found in the preSMA ROI (Study 1: mean beta = 0.083 ± 0.16 (SD), T_29_=2.78, P=0.0047; Study 2: mean beta = 0.057 ± 0.15 (SD), T_29_=2.14, P=0.021; **Fig. 8D-E**), consistent with the motor component of action imitation. In addition, an exploratory conjunction analysis showed that this imitation signal was tracked in a network of regions involved in action observation and action preparation, including the preSMA and SMA, but also the bilateral inferior parietal lobule, left motor cortex, and left dlPFC (**Fig. 8F**; see **Table S4** for details).

An interesting observation is that the preSMA ROI was found to represent both emulation and imitation update signals. Even though the two regressors were somewhat correlated (mean correlation coefficient R=0.347), they were included together as parametric modulators of the BOLD signal at the time of feedback and allowed to compete for variance. This suggests that activity in the preSMA uniquely contributes to each update process. In addition, we tested for a possible overlap between the two signals by extracting the corresponding clusters from the conjunction analysis. Interestingly, we found no overlap between the two clusters (**Fig. 8G**). The action change imitation cluster was exclusively located in the preSMA/SMA, while the token KL divergence emulation cluster was more frontal and anterior, extending into the dmPFC. This is consistent with a functional dissociation between the two update processes.

Other signals tested in this new SPM GLM3 were also found to be significant across studies (**Fig. S6, Table S4**). For example, during initial slot machine presentation on both observe and play trials, a vast network of regions was recruited when the partner’s most recent action was no longer available on the current trial. These included the dmPFC and preSMA ROIs (**Fig. S6A**), as well as the anterior insula, IFG, caudate nucleus, occipital and parietal regions (**Table S4**). At the time of choice on play trials, the propensity to choose according to imitation was negatively associated with preSMA activity (**Fig. S6B**), and the propensity to choose according to emulation was positively associated with mOFC activity (**Fig. S6C**). The mOFC and ACC were also found to track the value of the token obtained at the end of each trial (**Fig S6D**), consistent with the typical neural signature of value.

## Discussion

Across two independent fMRI studies, we provide neuro-computational evidence for an arbitration process between two observational learning strategies: imitation and emulation. We find that depending on the conditions of the environment, people flexibly adapt which strategy they preferentially rely on. Behavior was best explained by a computational model in which choice is a hybrid combination of imitation choice propensity and emulation choice propensity, weighted by a controller driven by the reliability of emulation. Using model comparison, we show that this arbitration model performed better than a model implementing each strategy individually. To our knowledge, these findings represent the first time an arbitration process between two learning strategies has been reported in the observational domain, in which instead of learning from experiencing outcomes, people learn from observing another agent.

Our fMRI results show that learning signals associated with each strategy were represented in distinct brain networks, when feedback is provided (i.e. during observation of the other agent’s action). When the observed agent chose a different action than on the previous trial, activity in the premotor cortex and inferior parietal cortex increased, possibly reflecting an update in the now preferred action according to imitation. Interestingly, this pattern of activity substantially overlaps with regions of the human mirror neuron system ^8,10,12^, consistent with the assumption that imitation learning relies on observing an action and repeating that same action in the future. The update in token values, calculated as the KL divergence between prior and posterior values, was reflected in a network of regions including the dmPFC, bilateral insula, right TPJ, IFG and dorsal striatum. Some of these regions (dmPFC and right TPJ) likely reflect an involvement of mentalizing abilities, in which the other agent’s goal has to be represented ^16^. The involvement of additional regions such as the dorsal striatum and IFG is consistent with previous literature on social learning, which has implicated these regions in inverse reinforcement learning ^15^, learning about another agent’s expertise ^20^, or tracking vicarious reward prediction errors ^34^. The IFG and anterior insula have also been found to play a role in attentional and executive control ^42,43^ and may thus reflect the engagement of emulation as a more complex strategy requiring increased cognitive and attentional resources. The fact that those distinct signals were identified at the same time point in the task suggests that the brain is keeping track of the decision values associated with each strategy in parallel, allowing individuals to deploy either strategy when needed.

The arbitration process, which in our final revised model is proposed to be driven exclusively by trial-by-trial variations in the reliability of emulation, was found to be correlated with BOLD responses in the right vlPFC, ACC, and bilateral TPJ. These findings suggest that these regions may be involved in implementing the arbitration process between the two OL strategies. Specifically, they may act as a hub in which information relevant to imitation learning (e.g. from premotor or inferior parietal cortex) and information relevant to emulation learning (e.g. from dmPFC or IFG) are dynamically integrated. Further evidence will need to be garnered for this possibility in follow-up work. For instance, establishing the nature of the effective connectivity between the arbitration regions and the brain regions involved in emulation and imitation respectively would add insight into how the arbitration process might be implemented at the network level. While beyond the scope of the present paper, this additional work can be conducted using the same datasets acquired here. Furthermore, future work could help establish the causal relevance of the structures identified in the arbitration process stimulating or inhibiting activity in those structures and establishing the effects on behavioral markers of emulation and imitation. This research agenda on OL, parallels a similar series of studies that has been conducted in the experiential domain, to explore the causal role of brain structures in arbitrating between model-based and model-free reinforcement-learning in the experiential domain ^27,44^. Interestingly, the right vlPFC region identified here was also found to track reliability signals related to arbitration between model-based (MB) and model-free (MF) RL in the experiential domain ^27^. However, in addition to the vlPFC regions identified in MB vs MF RL arbitration, in the present study we found evidence for other brain regions associated with the OL arbitration process not implicated in experiential arbitration, including the bilateral TPJ. The involvement of the TPJ, could relate to the fact that in order to compute the reliability of the emulation system, it is necessary to rely on computational mechanisms related to inference about an agent’s goals or intentions.

### A general arbitration mechanism for assigning control over behavior to different systems?

When compared with previous studies on the arbitration over MB and MF RL, our findings hint at important generalities about how control over different systems might be implemented in the brain. In both experiential and observational domains, it seems that uncertainty or reliability in the predictions of the systems might be a general mechanism for implementing arbitration between learning systems. Evidence for the use of reliability as a meta-control variable in at least two different behavioral contexts could impose constraints on general theoretical implementations of meta-control. If general purpose theories of meta-control assume arbitration is based on considerations about the expected value of particular control strategies and/or estimates of potential cognitive costs, they will have to take into consideration accumulating evidence that uncertainty and/or reliability of particular cognitive strategies is utilized at the computational and neural level to drive meta-control. That said, reliability of predictions need not be the only variable utilized to drive arbitration, and indeed in a recent study we found that task complexity is also an important variable in modulating the degree of engagement of MB and MF strategies in experiential learning ^45^. We also note that in the present study, the emulation reliability signal is minimal (and uncertainty maximal) when it is most difficult to compute which token is currently valuable from inspecting the token distributions on the slot machines. Thus, in the present case, cognitive difficulty is aligned with emulation (un)reliability.

At the neural level, the engagement of vlPFC in tracking both emulation reliability in the present study and MB / MF reliability in a past study ^27^, suggests at least some degree of overlap in the neural mechanisms of arbitration during observational and experiential learning. This could in turn suggest that the vlPFC plays a much more general role in arbitration between different strategies. One speculative idea is that this region could play a generic role in arbitrating between different strategies across multiple cognitive domains, as well as within cognitive domains. So, perhaps this same region might govern interactions between Pavlovian and instrumental control systems, as well as competition or cooperation between observational and experiential learning (should the two learning modalities provide different sources of information) as well as in mediating arbitration between different experiential and observational learning systems. Furthermore, it seems reasonable to postulate that arbitration doesn’t necessarily always have to involve competition between only two candidate systems, rather such competition might (subject to cognitive and attentional constraints), involve more than two systems. For instance, in observational learning, competition and/or cooperation could arise between emulation, imitation and (though not studied here) vicarious reinforcement-learning systems. Perhaps such three-way interactions might also be mediated by the same arbitration circuitry. Such an arbitration process could be generalized further to the selection between a multiplicity of strategies for behavioral control such as between many-fold model-based strategies that depend on different assumptions about state-space structure in the experiential domain ^46^. Future work will be needed to address all of these exciting possibilities.

### Distinct representations between imitation/emulation and model-based/model-free learning

One potential argument that could be made about the present findings is that we are merely recapitulating previous findings on the representation of MB and MF RL and the arbitration between them. In other words, could the computational processes we are labeling as emulation and imitation be mere implementations of MB and MF RL in an observational learning situation? We believe this argument does not hold water for the following reasons. First, by design, we have excluded the possibility that a simple extension of MF RL into the observational domain could explain the findings. In vicarious RL, an observer can co-opt the rewards experienced by another agent as if she has experienced them herself, and subsequently deploy a model-free learning strategy to acquire vicarious reward values for actions or stimuli ^6,34^. However, because in the present study we do not reveal the current reward value of tokens to our participant observers, a vicarious reward learning strategy cannot succeed in this situation. Vicarious RL is the closest to MF RL in the experiential literature and has been ruled out in the present case. Instead, the imitation strategy that we ultimately found to provide the best explanation for participant’s behavioral and neural data involves copying the action that the participant last saw the agent perform. This is distinct to MF RL, in that it involves learning about actions rather than rewards, and it does not include value computation over a history of recent events, simply a repeat of the most recent available action. Second, MB RL in the experiential domain is not typically assumed to involve the capacity for reverse inference – inferring the hidden goals of an agent based on observing that agent’s behavior in combination with observable information about the task structure ^15^. Instead, this type of inference is often described as “inverse” reinforcement learning in the machine learning literature ^47^, and constitutes a distinct class of algorithms to that of MB RL. Third, at the neural level, regions of the brain known to implement “mentalizing” computations, such as the TPJ, have been consistently implicated, in the present study and others ^15,18,20,21^, in tracking computations associated with the emulation system and/or the arbitration process. Such brain regions are not typically identified in studies of MB RL, suggesting that at the neural circuit level there is at least a partial distinction between them. Finally, brain regions implicated in action imitation in the present study (premotor and inferior parietal cortex) also do not cleanly map onto the areas involved in previous studies of reinforcement-learning whether MF or MB. Taken together, at the computational and neural levels, our findings suggest that we are not merely recapitulating the MB vs MF distinction.

### Replicability of computational neuroimaging findings

Beyond shedding light on the implementation of arbitration in observational learning, the present study is noteworthy for methodological reasons. Although replication is often recognized as the bed-rock for validation of scientific claims ^38,39,41^, inclusion of within-paper replications of fMRI studies is rare, perhaps because of how expensive data collection is for fMRI. Furthermore, for both computational model fitting and perhaps even more so for fMRI data analysis, the combination of a very large amount of data along with very high flexibility in data analysis pipelines, has been suggested to lead to an increased risk that reported findings may often be invalidated by modeler and/or experimenter degrees of freedom ^28–30^. Simply put, if one runs multiple analysis pipelines and focus on the ones that give the “best” results, the risk is that those results may depend on overfitting to noise in the data, as opposed to actual signal. Here we addressed this overfitting concern by implementing a replication study in which we pre-registered the full computational model specification, model-fitting and analysis pipeline for both the behavioral and fMRI data. Thus, we obtained a fully independent out-of-sample validation of our findings from the first study. We consider it to be an encouraging sign for typical computational behavior and fMRI studies that many of our initial findings were well replicated in the second study. Our computational model fitting results were closely replicated. In addition, a substantial subset of our fMRI results were replicated, especially those pertaining to the emulation system and the arbitration process, suggesting that some fMRI findings can be replicated even when analytical flexibility is virtually eliminated. It is also worth noting that our MRI scanner was upgraded from a Siemens Trio to Siemens Prisma platform between the two studies. Despite that substantial change in scanner hardware, brain activity patterns for most contrasts of interest were highly similar. Naturally we did try to keep the MR sequences and acquisition parameters we used as similar as possible across platforms, which of course is easier to do given the two scanners do come from the same manufacturer and do have many similarities. However, once one does keep flexibility in the analysis pipeline to a minimum, our findings are encouraging for the replicability of fMRI studies even across platforms.

### Exploratory analyses: using the quality of evidence from fMRI data to inform behavioral and computational processes

Intriguingly, those results that did not replicate well in the 2^nd^ fMRI dataset pertained to the imitation system, as implemented via our proposed action-based reinforcement-learning model. This subset of poorly replicated fMRI findings motivated us to revisit our proposed computational model for imitation. We then implemented a much simpler imitation strategy in which instead of keeping track of action prediction errors, the imitation system merely keeps track of which action the agent chose on the last occasion that action was available. Furthermore, our fMRI data also suggested that the arbitration process should be modified as we found no evidence that survived replication to suggest that imitation reliability was tracked in the brain. Indeed, the reliability difference signal we proposed to mediate arbitration in Study 1 was only really tracking emulation reliability and not imitation reliability when subjected to closer inspection. Thus, we implemented a new arbitration scheme that assigned control to emulation or imitation based on emulation reliability only. Our new combined model with revised imitation and arbitration mechanisms was found to clearly outperform our original model in terms of fits to both behavior and BOLD responses. Moreover, this revised model also produced findings about the neural representation of imitation and the arbitration process that were more clearly replicated across studies. Thus, we used knowledge about the quality of the evidence we gleaned from the brain to revisit our original hypotheses, which is a nice example of how evidence from neural data can be used to inform computational and psychological theory. While these additional analyses were not part of the pre-registered confirmatory process and must thus be labeled as exploratory, we strongly believe they are likely to reveal robust mechanisms because they generalize across two separate datasets, and there is a direct link between the robustness of the behavioral model fits and robustness of the fMRI results. That said, subsequent work will ideally subject this new model and findings to further confirmatory replication.

### General conclusion

Imitation and emulation have been studied at length in comparative and developmental psychology studies ^3,48–50^ and thus are of significance for many fields of research, from education to evolutionary psychology. Here we developed and optimized a novel paradigm and associated neuro-computational modelling approach that allowed us to adequately separate the mechanisms of imitation and emulation as observational learning strategies. We illuminate the behavioral and neural signature of how these two strategies compete for control over behavior in a reliability-driven arbitration process.

## Materials and Methods

### Participants

Thirty healthy participants (12 females, 18 males, mean age = 31.67 ± 4.94 (SD)) took part in Study 1 between November 2017 and January 2018. For the replication study (Study 2), 33 healthy participants were recruited between October 2018 and January 2019. Three participants were excluded for excessive head motion in the scanner (N=1), incidental finding (N=1) and missing more than 20% of responses on the task (N=1). As preregistered, our final sample for Study 2 included 30 participants (12 females, 18 males, mean age = 31.2 ± 8.15 (SD)). There was no age (t_58_=0.27, p=0.79) or gender difference across studies. All participants met MRI safety criteria, had normal or corrected-to-normal vision, no psychiatric/neurological conditions, and were free of drugs for 7 days prior to the scan that might potentially interfere with the BOLD response (cannabis, hallucinogenic drugs). They were paid $20 per hour, in addition to bonus money earned during the task ($5 to $8) depending on their performance. The research was approved by the Caltech Institutional Review Board, and all participants provided informed consent prior to their participation.

### Experimental design

Participants performed a task in which they have to choose between slot machines in order to maximize their chances of winning a valuable token (worth $0.10). They were instructed that there are 3 tokens available in the game (green, red, or blue) and that at any given time, only one token is valuable and the other two are worth nothing. When arriving to the lab, participants first completed an experiential version of the task (∼5 minutes) in which the computer told them at the beginning of each trials which token is valuable. They were then presented with the 3 slot machines and instructed that the proportion of green, blue and red colors on each slot machine corresponds to the probability of obtaining each token upon choosing that slot machine. In addition, on each trial one of the slot machines was greyed out an unavailable; therefore, participants had to choose between the remaining two active slot machines.

During the main task (observational learning, **Fig. 1A**), participants were instructed that the valuable token would switch many times during the task, but they would not be told when the switches occur anymore. Instead, they would have to rely on observing the performance of another agent playing the task. On 2/3 of trials (‘observe’ trials), participants observed that other agent play and knew that this agent had full information about the valuable token and was therefore performing 100% correctly. On 1/3 of trials (‘play’ trials), participants played for themselves and the sum of all play trial outcomes was added to their final bonus payment.

Participants completed a practice of the observational learning task before scanning (2 blocks of 30 trials), followed by 8 blocks of 30 trials of the task while undergoing fMRI scanning. Each block of 30 trials contained 20 observe trials and 10 play trials. The sequence of trials within each block was pre-determined with simulation in order to maximize learning. Block order was counterbalanced across subjects.

Four conditions were implemented in a 2 (stable vs volatile) by 2 (low vs high slot machine uncertainty) design across blocks. Volatility was manipulated by changing the frequency of token switches (**Fig. 1B**): there was one switch in the valuable token during stable blocks, and 5 switches during volatile blocks. Uncertainty associated with the slot machines was experimentally manipulated by changing the token probability distribution associated with each slot machine (**Fig. 1C**): [0.75, 0.2, 0.05] in low uncertainty blocks and [0.5, 0.3, 0.2] in high uncertainty blocks.

Trial timings are depicted in **Fig. 1A**. Trial type (“Observe” or “Play” printed on the screen) was displayed for 1s, immediately followed by the presentation of the slot machine for 2s. On observe trials, there was then a jittered fixation cross (1-4s), followed by the video showing the choice of the partner (around 2s). After another jittered fixation cross (1-4s), the token obtained by the partner was shown on screen for 1s. On play trials, the slot machine presentation was immediately followed by the onset of the word “CHOOSE” indicating participants they had 2s to make their choice. The chosen slot machine was highlighted for 0.5s, followed by a jittered fixation cross (1-4s) and the presentation of the token obtained by the participant. Finally, there was a jittered inter-trial interval of 1-5s.

The procedure and task was exactly the same between Study 1 and Study 2.

### Behavioral analysis

To test for the presence of the two learning strategies (imitation, based on learning from previous partner’s actions, versus emulation, based on learning from previous evidence about valuable token), a general linear model (GLM) was run using the glmfit function on Matlab. Specifically, the dependent variable was choice of left (coded as 1) or right (coded as slot machine, and the two independent variables (regressors) were constructed as follows:

- Effect of past actions: for each previous observe trial between last switch in valuable token and current play trial, the other agent’s action was coded as +1 if the current left-most slot machine was chosen, −1 if it was unchosen or 0 if it was unavailable. The value of the regressor at each play trial was calculated as the sum of these past actions scores, which represents the accumulated evidence for the left slot machine given past actions chosen by the other agent.
- Effect of past tokens: for each previous observe trial between last switch in valuable token and current play trial, token information can be inferred (e.g. “green is the valuable token for sure”, or “the valuable token could be green or blue”). From this, the probability that the left (vs right) slot machine results in the valuable token was calculated based on token probability distribution associated with each slot machine. The value of the regressor at each play trial was calculated as the sum of these probability differences, which represents accumulated evidence for the left slot machine given past token information.

We ran this GLM for each participant, averaged the resulting beta values across all participants and tested their significance with permutation tests (10,000 permutations), since data were usually not normally distributed across the sample.

### Computational models of behavior

As reported in the preregistration, a total of 9 computational models of behavior were tested, split into 5 classes of models.

#### 1) Approximate Bayesian Emulation Models

In these models, emulation learning is based on a multiplicative update of the probability of each token being valuable, *V*_*g*_, *V*_*r*_, and *V*_*b*_, for green, red, and blue tokens respectively. At t=0, all values are initialized at 1/3. The update occurs after observing the partner’s action (example for green token):

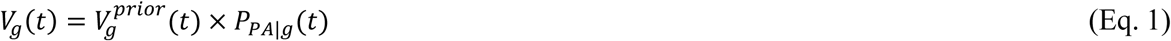

where *P*_*PA*|*g*_(*t*) is the probability of observing the partner’s action given that green is the valuable token on trial t. Given that the partner is always correct, *P*_*PA*|*g*_(*t*) equals either 1 or 0.

The prior value is calculated as follows:

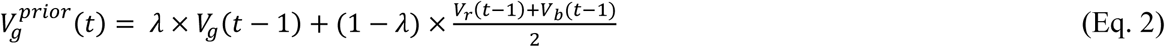

The parameter λ represents trust in current estimates of token values and allows for switches to happen by resetting reward probability of each token to a non-zero value on each trial. Our simulations showed that the value of this parameter that maximizes model’s inference performance changes when volatility is high (token switch frequency higher than 0.2, i.e. more than one switch every 5 trials). However, in our task, token switch frequency is 0.067 per trial for stable blocks and 0.167 per trial for volatile blocks. In these conditions, the value of λ that maximizes performance is as close as possible to 1, but with a small leak to allow the values to be updated on the next trials. Therefore, we used λ=0.99. However, if participants overestimate volatility in the environment, a model with a smaller λ could capture behavior of such participants better. We thus tested two models: one with a fixed λ of 0.99 (**Model 1**) and one allowing the λ parameter to vary for each participant (**Model 2**).

Token values *V*_*g*_, *V*_*r*_, and *V*_*b*_ are then normalized so that they sum to 1 ^51^. Then the value of choosing each slot machine 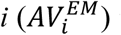 is computed through a linear combination of token values and token probabilities (*p*_*g*_, *p*_*r*_, *p*_*b*_) given by the slot machine:

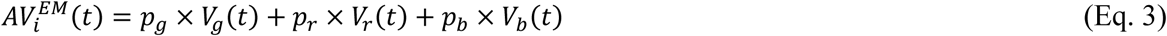

Finally, decision value is calculated as a soft-max function of the difference in value between the two available slot machines on the current play trial.

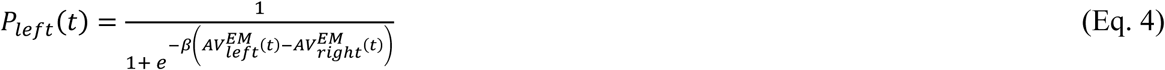

where β is an inverse temperature parameter, estimated for each subject.

#### 2) Action Imitation RL Models

These models were implemented as reinforcement learning (RL) models in which the value of each action is updated after every observation depending on whether it was chosen by the partner or not. On the first trial, action values (AV) are initialized at 0. Actions chosen by the other agent are updated positively, while unchosen actions are updated negatively, both according to a learning rate α:

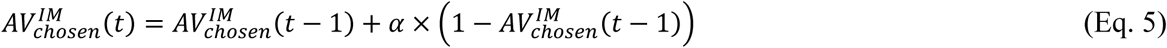

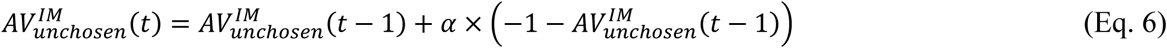

The learning rate α was either estimated as a fixed parameter for each subject (**Model 3**) or varied over time depending on recency-weighted accumulation of unsigned prediction errors with weight parameter η and initial learning rate α_0_ (**Model 4**) ^52,53^:

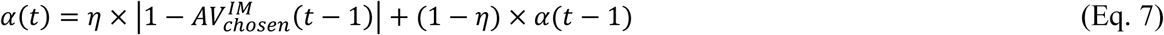

Decision rule is implemented using Eq. 4.

#### 3) Emulation RL Models

Two additional models were defined to test the possibility that emulation is implemented as an RL process, rather than as a multiplicative update as described in Models 1 and 2. The value of each token (example below for the green token g) is initialized as 0, and updated based on a token prediction error (TPE) and a learning rate α:

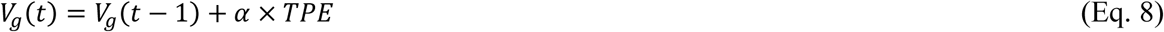

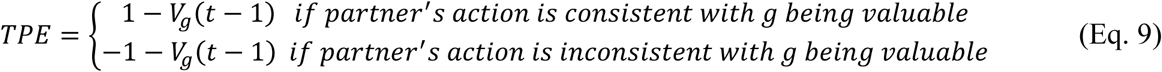

Similarly to the imitation models above, the learning rate α was either estimated as a fixed parameter for each subject (**Model 5**) or varied over time depending on recency-weighted accumulation of unsigned prediction errors (|*TPE*|) with weight parameter η and initial learning rate α_0_ (**Model 6**).

Action values are then calculated from token values using Eq. 3, and decision rule is implemented using Eq. 4.

#### 4) Arbitration Models

Arbitration was governed by the relative reliability of emulation (R^EM^) and imitation (R^IM^) strategies. R^EM^ is driven by the min-max normalized Shannon entropy of emulation action values (i.e. the slot machines action values AV_i_ predicted by the Approximate Bayesian Emulation Models described above):

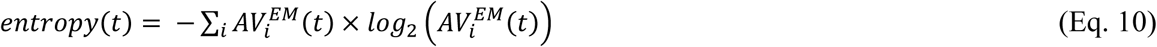

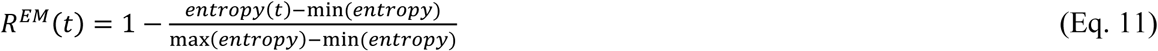

R^IM^ is driven by the min-max normalized unsigned action prediction error (APE):

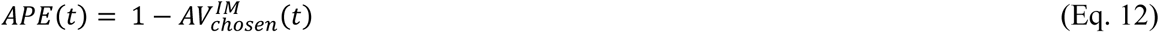

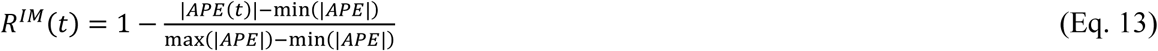

Minimum and maximum entropy and |APE| values were obtained from practice trial data, by fitting the emulation-only and imitation-only model to that practice data and extracting the minimum and maximum values from the two variables.

This definition of reliability suggests that when entropy between slot machines is high (driven both by uncertainty about which token is valuable and by the uncertainty manipulation depicted in **Fig. 1C**), emulation becomes unreliable. When action prediction errors are high (driven by unexpected partner’s actions), imitation becomes unreliable.

Arbitration is then governed by an arbitration weight ω, implemented as a soft-max function of the reliability difference, with the addition of a bias parameter δ(δ>0 reflects a bias towards emulation, δ<0 reflects a bias towards imitation):

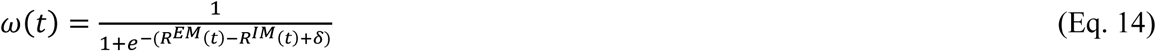

The probability of choosing the slot machine on the left is computed separately for each strategy:

- using Eqs. 1-4 for emulation 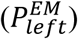, with either an optimal λ of 0.99 (**Model 7**) or an estimated λ parameter for each subject (**Model 8**), and with inverse temperature parameter *β*^*EM*^
- using Eqs. 5, 6 and 4 for imitation 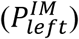, with fixed learning rate α and with inverse temperature parameter *β*^*IM*^

Then the two decision values 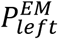 and 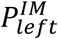 are combined using the arbitration weight ω:

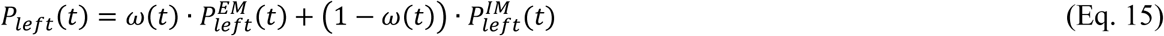

#### 5) Outcome RL Model

One last model we tested (**Model 9**) is the possibility that participants mistakenly learn from the token that is presented as an outcome at the end of the trial, instead of learning from the partner’s actions. This was implemented similarly to other RL models. The value of each token was updated positively every time that token was obtained as an outcome, either by the partner or by the participant, and negatively if that token was not obtained. Action values and decision value were then calculated using Eqs. 3 and 4.

#### 6) Exploratory Arbitration Model

This model (**Model 10**) was defined to test the possibility that imitation is implemented as a simpler 1-step learning strategy in which the most recent partner’s action is repeated on the current trial. Specifically, the probability of choosing the left slot machine on each play trials according to imitation is calculated as follows:

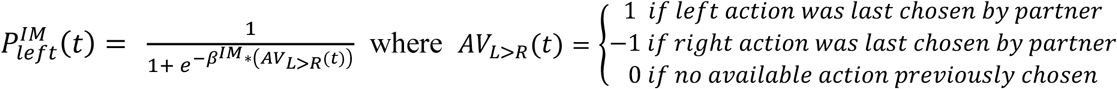

The probability of choosing left according to emulation 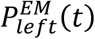 was defined as above (Eqs. 1-4). Arbitration in this model was driven exclusively by the reliability of the emulation strategy (Eq. 11), with the arbitration weight ω calculated as a soft-max function of the emulation reliability, with the addition of a bias parameter δ. Then the two decision values 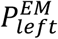 and 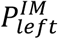 are then combined with arbitration weight ω like above (Eq. 15).

### Model fitting and comparison

As preregistered, model fitting and comparison were performed in two different ways to assess robustness of model-fitting results:

1. Using maximum likelihood estimation in Matlab with the fminunc function to estimate parameter estimated for each subject, followed by an out-of-sample predictive accuracy calculation to compare models. Specifically, for the accuracy calculation, subjects were split into 5 groups of 6 subjects, mean parameters were estimated for 4 groups (24 subjects) and tested on the remaining group (6 subjects). This was repeated for all groups, as well as for 100 different groupings of subjects. Mean predictive accuracy (proportion of subjects’ choices correctly predicted by the model) for each model is reported in **Table 1**.
2. Using hierarchical Bayesian random effects analysis. Following ^35,36^, the (suitably transformed) parameters of each participant are treated as a random sample from a Gaussian distribution characterizing the population. We estimated the mean and variance of the distribution by an Expectation-Maximization method with a Laplace approximation. We estimated each model’s parameters using this procedure, and then compared the goodness of fit for the different models according to their group-level integrated Bayesian Information Criteria (iBIC, see **Table 1**). The iBIC was computed by integrating out individual subjects’ parameters through sampling. The full method is described in e.g. ^35,36^.

### Posterior predictive analysis

We tested that the winning model could reliably predict the behavioral effects obtained by simple GLM (see “***Behavioral analysis***” paragraph above), namely the effects of past actions and past tokens on current choice. To do so, we used individual subject’s parameters from the winning model (**Model 7**), as well as individual subject’s parameters from the simple emulation (**Model 2**) and imitation (**Model 3**) models, to generate hypothetical choice data for each participant using that participant’s trial sequence. We then ran the same GLM on the model-generated data for each participant, calculated the mean GLM betas across participants and repeated the process (data generation + GLM fitting) 1000 times. The GLM betas from these 1000 iterations are plotted as histograms on **Fig. 2C-D**, together with the true effect on participants’ actual behavioral data (red point ± standard error).

We also examined our prediction that the use of imitation versus emulation strategies is modulated by action volatility and by token uncertainty. To do so, we extracted the arbitration weight ω(t) from **Model 7** for each trial and each participant, and averaged it for each of the 4 conditions (**Fig. 3A-B**): stable/low uncertainty, volatile/low uncertainty, stable/high uncertainty, volatile/high uncertainty. Differences in arbitration weight across conditions were tested in a 2-by-2 repeated-measures ANOVA. In a separate analysis, we compared the mean likelihood per trial of imitation (**Model 3**) and emulation (**Model 2**) for each of the 4 conditions (**Fig. 3C-D**).

### Software

The task was coded and presented using PsychoPy ^54^ version 1.85 under Windows. Behavioral analyses, including computational models, were run on Matlab (R2018a). MRI data was analyzed using FMRIB Software Library (FSL; version 5.0; https://fsl.fmrib.ox.ac.uk/fsl/fslwiki), Advanced Normalization Tools (ANTs; http://stnava.github.io/ANTs/) and Statistical Parametric Mapping (SPM; version 12; https://www.fil.ion.ucl.ac.uk/spm/software/spm12/).

### fMRI data acquisition

For Study 1, fMRI data was acquired on a Siemens Magneto TrioTim 3T scanner at the Caltech Brain Imaging Center (Pasadena, CA), which was later upgraded to a Siemens Prisma 3T scanner before Study 2. The same 32-channel radio frequency coil was used for both studies. MRI acquisition protocols and sequences were also kept as similar as possible. For functional runs, 8 scans of 410 volumes each were collected using a multi-band echo-planar imaging (EPI) sequence with the following parameters: 56 axial slices (whole-brain), A-P phase encoding, −30 degrees slice orientation from AC-PC line, echo time (TE) of 30ms, multi-band acceleration of 4, repetition time (TR) of 1000ms, 60-degree flip angle, 2.5mm isotropic resolution, 200mm x 200mm field of view, EPI factor of 80, echo spacing of 0.54ms. Positive and negative polarity EPI-based fieldmaps were collected before each block with very similar factors as the functional sequence described above (same acquisition box, number of slices, resolution, echo spacing, bandwidtch and EPI factor), single band, TE of 50ms, TR of 4800ms (Study 1)/ 4810ms (Study 2), 90-degree flip angle. T1-weighted and T2-weighted scans were also acquired either at the end of the session or halfway through, both with sagittal orientation, field of view of 256mm x 256mm, and 1mm (Study 1)/0.9mm (Study 2) isotropic resolution.

### fMRI data preprocessing

The same preprocessing pipeline was used in both studies. First, reorientation and rough brain extraction of all scans were performed using fslreorient2std and bet FSL commands, respectively. Fieldmaps were extracted using FSL topup. Following alignment of the T2 to the T1 (FSL flirt command), T1 and T2 were co-registered into standard space using ANTs (CIT168 high resolution T1 and T2 templates ^55^). Then an independent component analysis (ICA) was performed on all functional scans using FSL MELODIC; components were classified as signal or noise using a classifier that was trained on previous datasets from the lab; and noise components were removed from the signal using FSL fix. De-noised functional scans were then unwarped with fieldmaps using FSL fugue, co-registered into standard space using ANTs and skull-stripped using SPM imcalc. Finally, 6mm full-width at half maximum Gaussian smoothing was performed using SPM.

### fMRI data modelling – preregistered

Two separate GLMs were used to model the BOLD signal, incorporating an AR(1) model of serial correlations and a high-pass filter at 128Hz.

#### SPM GLM1

The first GLM was built to examine the neural correlates of arbitration and included the following regressors:

- Slot machine onset – observe trials (1), parametrically modulated by (2) the difference in reliability (*R*^*EM*^ − *R*^*IM*^), (3) the difference in available action values as predicted by the imitation strategy 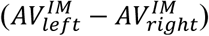, and (4) the entropy over the 3 token values as predicted by the emulation strategy (− ∑_*token*_ *V* · *log*_2_ *V*).
- Slot machine onset – play trials (5), parametrically modulated by (6) the difference in reliability (*R*^*EM*^ − *R*^*IM*^), and (7) the chosen action value (expected reward probability) as predicted by the arbitration model.
- Partner’s action onset – observe trials (8), parametrically modulated by (9) the difference in reliability (*R*^*EM*^ − *R*^*IM*^), and (10) the reduction in entropy over token values calculated as the KL divergence between prior and posterior token values predicted by the arbitration model.
- Token onset – observe trials (11), parametrically modulated by (12) the difference in reliability (*R*^*EM*^ − *R*^*IM*^), and (13) an observational reward prediction error (oRPE), calculated as the difference between the initial expected reward value given the chosen slot machine, and the posterior value of the token shown on screen (as predicted by the model).
- Token onset – play trials (14), parametrically modulated by (15) an experiential reward prediction error (eRPE), calculated as described above. We hypothesized that the difference in reliability would not occur at this onset given that it is not associated with any learning (occurring only during observe trials) or choice (occurring earlier during play trials).

#### SPM GLM2

The second GLM was identical to the first except that each arbitration-related regressor was separated into its emulation (*R*^*EM*^) and imitation (*R*^*IM*^) components. In addition, the chosen action value regressor used during play trials slot machine onset was replaced by two action value difference (chosen vs unchosen) regressors predicted by the imitation and emulation strategy separately. This model allowed looking for a neural signature of each strategy.

For both GLMs, trials from all 8 blocks were collapsed into one session in the design matrix. Regressors of no interest included missed choice onsets (if any) as well as 7 regressors modelling the transitions between blocks. All onsets were modeled as stick functions (duration = 0 s). All parametric modulators associated with the same onset regressors were allowed to compete for variance (no serial orthogonalization). GLMs were estimated using SPM’s canonical HRF only (no derivatives) and SPM’s classical method (restricted maximum likelihood).

First-level contrast images were created through a linear combination of the resulting beta images. For the reliability difference signal (SPM GLM1) and for individual reliability signals (SPM GLM2), first-level contrasts were defined as the sum of the corresponding beta images across all onsets where the parametric modulator was added. A global RPE signal was also examined by summing over the oRPE and eRPE contrasts.

### fMRI data modelling – exploratory

A third fMRI model, SPM GLM3, was defined to test the neural implementation of the behavioral arbitration **Model 10**, in which imitation is implemented as a simpler 1-step learning strategy of repeating the partner’s most recent action. The regressors were as follows:

- Slot machine onset – observe trials (1), parametrically modulated by (2) the reliability of emulation (*R*^*EM*^), and (3) whether the partner’s previous action is available or not.
- Slot machine onset – play trials (4), parametrically modulated by (5) the reliability of emulation (*R*^*EM*^), (6) whether the partner’s previous action is available or not, and the propensity to choose according to imitation (7) or according to emulation (8), as predicted by the arbitration model.
- Partner’s action onset – observe trials (9), parametrically modulated by (10) the reliability of emulation (*R*^*EM*^), (11) the KL divergence between prior and posterior token values predicted by the arbitration model, and (12) whether the partner’s most recent action is repeated, not repeated or unavailable on the current trial, coded as 1, −1 and 0 respectively.
- Token onset – observe trials (13), parametrically modulated by (14) the reliability of emulation (*R*^*EM*^), and (15) the value of the token shown on screen (as predicted by the model).
- Token onset – play trials (16), parametrically modulated by (17) the value of the token shown on screen (as predicted by the model).

### Regions of interest

Based on previous literature on observational learning ^15^ and arbitration processes between learning strategies ^27^, as well as the Neurosynth (http://neurosynth.org/) “Theory of Mind” meta-analysis map, the following 8 ROIs were defined and pre-registered:

- Left and right TPJ/pSTS: two 8-mm radius spheres around peaks of the Neurosynth map: (−54,-53,22) and (58,-58,20) for left and right, respectively.
- Medial OFC: 8-mm radius sphere around peak of the Neurosynth map (2,49,-20).
- dmPFC: 8-mm radius sphere around peak activation tracking expected value in other-referential space ^15^: (0,40,40).
- Pre-SMA/dACC: 8-mm radius sphere around peak activation tracking entropy reduction (KL divergence, ^15^: (−6,18,44).
- Left and right vlPFC: two 8-mm radius spheres around peak activations tracking maximum reliability of model-free and model-based learning ^27^: (−54,38,3) and (48,35,-2) for left and right, respectively.
- Dorsal striatum: anatomical bilateral caudate mask from AAL atlas ^56^.

Parameter estimates from the different contrasts were extracted from the ROIs and averaged across voxels within the ROI.

### fMRI model comparison

Model comparison and selection between SPM GLMs was performed using the MACS (Model Assessment, Comparison and Selection) toolbox for SPM ^57^ and included the following steps. For each subject and each model, cross-validated log model evidence (cvLME) maps were estimated. cvLME maps rely on Bayesian estimations of the models and Bayesian marginal likelihood to calculate, for each voxel, a voxel-wise cross-validated log model evidence for that model. Then, cross-validated Bayesian model selection ^58^ (cvBMS) was performed to compare GLMs. In cvBMS, second-level model inference is performed using random-effects Bayesian model selection, leading to voxel-wise model selection via exceedance probability maps. For each voxel in a grey matter mask, the exceedance probability was calculated as the posterior probability that a model is more frequent than any other model in the model space (**Fig. S5**). Exceedance probability was also averaged across voxels in the different ROIs (**Table S3**).

### Group-level inference and conjunction analysis

Second-level T-maps were constructed separately for each study by combining each subject’s first level contrasts with the standard summary statistics approach to random-effects analysis implemented in SPM. To assess the evidence for consistent effects across studies, conjunction maps were calculated with the minimum T-statistic approach ^59^ for each contrast of interest in the winning SPM GLM (**Table S4**), combining the second-level T-maps of each study. We thresholded conjunction maps at a conjunction P-value of P_conjunction_<0.0001 uncorrected, and minimum cluster size of 30 voxels, corresponding to a whole-brain cluster-level family-wise error corrected P-value of P_FWE_<0.05.

## Acknowledgements

This work was funded by the Caltech Conte Center for the Neurobiology of Social Decision-Making. K.I. is supported by the Japan Society for the Promotion of Science and the Swartz Foundation.

## Author contributions

C.C. and J.O. were responsible for conceptualization and methodology. C.C. carried out the investigations. C.C. and K.I. performed the analyses. C.C. and J.O. wrote the original manuscript draft. C.C., K.I. and J.O. reviewed and edited the manuscript. J.O. supervised the study and acquired funding.

