## Supplementary Materials for "Neuro-computational account of arbitration between imitation and emulation during human observational learning"

| | | Imitation signals | | Emulation signals | | | | Arbitration signals | |
|---|---|---|---|---|---|---|---|---|---|
| | | GLM2: Imitation reliability | GLM1: Imitation action value diff. | GLM2: Emulation reliability | GLM1: Token entropy | GLM1: Token KLdiv. | GLM1: Token RPE | GLM1: Reliability difference | GLM1: Chosen action value |
| Left TPJ pSTS | Study 1 | -0.046 ±0.46 | 0.088* ±0.17 | **0.172* ±0.55** | 0.083* ±0.16 | -0.010 ±0.17 | 0.042 ±0.14 | **0.201* ±0.53** | 0.032 ±0.23 |
| | Study 2 | - | -0.022 ±0.17 | **0.195* ±0.55** | -0.112* ±0.25 | - | - | **0.182* ±0.38** | - |
| Right TPJ pSTS | Study 1 | 0.041 ±0.65 | 0.088* ±0.21 | 0.299* ±0.66 | 0.121* ±0.26 | 0.046 ±0.21 | -0.032 ±0.23 | 0.277* ±0.59 | 0.022 ±0.24 |
| | Study 2 | - | -0.038 ±0.22 | 0.137 ±0.78 | -0.078 ±0.30 | - | - | 0.071 ±0.70 | - |
| mOFC | Study 1 | 0.387* ±0.58 | 0.020 ±0.15 | 0.128 ±0.86 | **-0.080* ±0.25** | -0.029 ±0.27 | 0.001 ±0.25 | -0.059 ±0.85 | **0.110* ±0.28** |
| | Study 2 | 0.077 ±0.28 | - | - | **-0.107* ±0.19** | - | - | - | **0.109* ±0.22** |
| dmPFC | Study 1 | -0.228 ±0.86 | 0.168* ±0.31 | 0.161 ±1.01 | 0.234* ±0.31 | **0.201* ±0.23** | 0.087 ±0.34 | **0.383* ±0.89** | -0.095 ±0.41 |
| | Study 2 | - | -0.018 ±0.20 | - | -0.061 ±0.27 | **0.098* ±0.21** | - | **0.228* ±0.54** | - |
| PreSMA dACC | Study 1 | -0.086 ±0.54 | 0.095* ±0.16 | 0.034 ±0.66 | 0.147* ±0.17 | **0.170* ±0.18** | 0.023 ±0.23 | 0.136 ±0.58 | **-0.144* ±0.29** |
| | Study 2 | - | 0.025 ±0.12 | - | -0.018 ±0.12 | **0.123* ±0.17** | - | - | **-0.135* ±0.22** |
| Left vlPFC | Study 1 | -0.007 ±0.78 | 0.130* ±0.27 | 0.187 ±0.76 | 0.133* ±0.26 | 0.066 ±0.24 | 0.116 ±0.29 | 0.209 ±0.68 | -0.084 ±0.40 |
| | Study 2 | - | 0.024 ±0.22 | - | -0.077* ±0.22 | - | - | - | - |
| Right vlPFC | Study 1 | 0.003 ±0.48 | 0.064* ±0.19 | **0.320* ±0.50** | 0.118* ±0.14 | 0.048 ±0.18 | 0.006 ±0.16 | 0.250* ±0.48 | 0.002 ±0.26 |
| | Study 2 | - | -0.041 ±0.14 | **0.186* ±0.39** | -0.045 ±0.18 | - | - | 0.035 ±0.43 | - |
| Dorsal striatum | Study 1 | -0.052 ±0.30 | 0.052* ±0.10 | -0.029 ±0.38 | 0.052* ±0.12 | **0.043* ±0.09** | 0.071* ±0.08 | 0.043 ±0.30 | -0.004 ±0.16 |
| | Study 2 | - | -0.013 ±0.068 | - | -0.022 ±0.08 | **0.033* ±0.09** | -0.030 ±0.13 | - | - |

**Table S1. Pre-registered ROI results.** Betas associated with the preregistered contrasts from SPM GLM 1 and 2 were extracted from the preregistered ROIs (average across all voxels in ROI) for each subject. This table reports the mean betas ± standard deviation across subjects, separately for each study. In Study 1, every contrast was examined in every ROI as a way to generate hypotheses. In Study 2, only significant effects from Study 1 were examined. * $P<0.05$, t-tests and permutation tests. Results highlighted in bold indicate replication (significant effects in the same direction) across studies.

| Region | Study 1 | | | | | Study 2 | | |
|---|---|---|---|---|---|---|---|---|
| | x | y | z | Cluster size | $T_{S1}$ | $Beta_{S2}$ (mean) | $SD_{S2}$ | $P_{S2}$ |
| *Imitation reliability (>0)* | | | | | | | | |
| mOFC, vmPFC, ACC | 3 | 37 | -7 | 325 | 5.01 | 0.038 | 0.35 | 0.28 |
| *Imitation reliability (<0)* | | | | | | | | |
| Right inferior Parietal / Angular gyrus | 48 | -46 | 58 | 107 | 4.43 | -0.065 | 0.59 | 0.27 |
| *Imitation action value difference (>0)* | | | | | | | | |
| Right dlPFC | 30 | 24 | 41 | 333 | 5.82 | -0.017 | 0.14 | 0.25 |
| Right TPJ | 48 | -49 | 36 | 333 | 5.39 | -0.016 | 0.30 | 0.39 |
| **Pre-SMA / dmPFC** | **-5** | **24** | **53** | **193** | **4.74** | **0.034** | **0.11** | **0.05** |
| Left dlPFC | -43 | 19 | 41 | 101 | 4.64 | 0.072 | 0.24 | 0.06 |
| Precuneus | 3 | -61 | 18 | 216 | 4.46 | -0.050 | 0.18 | 0.07 |
| Left Thalamus | -15 | -29 | 11 | 104 | 4.42 | -0.012 | 0.08 | 0.21 |
| Left TPJ | -48 | -59 | 36 | 275 | 4.28 | 0.008 | 0.32 | 0.45 |
| *Emulation reliability (>0)* | | | | | | | | |
| **Right anterior Insula** | **43** | **17** | **-12** | **126** | **4.82** | **0.258** | **0.43** | **<0.001** |
| *Token entropy (>0)* | | | | | | | | |
| Bilateral inferior Parietal / Angular gyrus / TPJ / Precuneus | 38 | -49 | 46 | 3755 | 7.40 | 0.023 | 0.28 | 0.33 |
| Right dlPFC / IFG / OFC / vlPFC | 28 | 9 | 61 | 2875 | 6.20 | -0.004 | 0.15 | 0.44 |
| Left dlPFC / IFG / OFC / vlPFC | -20 | -4 | 63 | 2485 | 6.08 | -0.024 | 0.16 | 0.20 |
| Cerebellum | -10 | -79 | -25 | 203 | 5.88 | -0.002 | 0.12 | 0.47 |
| Right mid-Temporal | 55 | -41 | -5 | 215 | 5.82 | 0.006 | 0.13 | 0.39 |
| dmPFC / Pre-SMA/ dACC | -8 | 29 | 41 | 708 | 5.31 | -0.029 | 0.17 | 0.18 |
| Thalamus | 0 | -16 | 3 | 191 | 4.95 | -0.003 | 0.09 | 0.44 |
| *Token KL divergence (>0)* | | | | | | | | |
| **Left anterior Insula** | **-33** | **14** | **-10** | **119** | **4.88** | **0.102** | **0.15** | **<0.001** |
| **Right IFG / Precentral gyrus** | **35** | **9** | **33** | **342** | **4.69** | **0.157** | **0.23** | **<0.001** |
| **Right anterior Insula** | **40** | **19** | **-2** | **122** | **4.65** | **0.117** | **0.15** | **<0.001** |
| **Pre-SMA / dACC** | **-8** | **19** | **46** | **140** | **4.44** | **0.174** | **0.22** | **<0.001** |
| **Left IFG / Precentral gyrus** | **-48** | **7** | **26** | **103** | **4.41** | **0.187** | **0.27** | **<0.001** |
| **Right Supramarginal / inferior Parietal** | **53** | **-39** | **46** | **109** | **4.09** | **0.246** | **0.38** | **<0.001** |
| *Reliability difference – EM vs IM* | | | | | | | | |
| **Right anterior Insula** | **40** | **17** | **-12** | **113** | **5.97** | **0.099** | **0.28** | **0.03** |
| **Right IFG** | **45** | **4** | **21** | **184** | **5.02** | **0.173** | **0.51** | **0.04** |
| **ACC / dmPFC** | **13** | **44** | **26** | **91** | **4.80** | **0.089** | **0.23** | **0.02** |
| **Right Angular gyrus** | **40** | **-74** | **48** | **206** | **4.38** | **0.225** | **0.74** | **0.05** |

**Table S2. Replication findings using clusters from Study 1 group-level maps as functional ROIs in Study 2.** Significant activation clusters from Study 1 (pre-registered) were identified following whole-brain cluster-level FWE correction at $p<0.05$ and cluster forming threshold at $P<0.001$ uncorrected. Peak MNI coordinates are reported for Study 1, together with cluster size (number of contiguous voxels in the cluster) and peak voxel T-value. Each significant cluster was saved as a functional ROI, and mean signal from Study 2 was extracted in this ROI and averaged across subjects to assess replication. Mean beta, standard deviation and p-value (t-tests, also confirmed with permutation tests) are reported for Study 2. Regions highlighted in bold indicate replication at $P\leq0.05$.

| | Study 1 | | Study 2 | |
|---|---|---|---|---|
| | SPM GLM2 | SPM GLM3 | SPM GLM2 | SPM GLM3 |
| Left TPJ pSTS | 0.366 | **0.634** | 0.028 | **0.972** |
| Right TPJ pSTS | 0.257 | **0.743** | 0.159 | **0.841** |
| mOFC | 0.047 | **0.953** | 0.013 | **0.987** |
| dmPFC | **0.519** | 0.481 | 0.031 | **0.969** |
| PreSMA dACC | 0.110 | **0.890** | 0.009 | **0.991** |
| Left vlPFC | 0.201 | **0.799** | 0.085 | **0.915** |
| Right vlPFC | 0.029 | **0.971** | 0.050 | **0.950** |
| Dorsal striatum | 0.035 | **0.965** | 0.019 | **0.981** |

**Table S3. Exceedance probabilities from Bayesian fMRI model selection in pre-registered ROIs.** After performing the Bayesian model selection analysis between SPM GLM2 and SPM GLM3, we averaged the exceedance probability associated with each model across all voxels of a given ROI. For each ROI, this exceedance probability represents the posterior probability that a model is more frequent than the other. For all ROIs and across both studies (except in the dmPFC in Study 1), SPM GLM3 was found to explain variations in the BOLD signal substantially better than SPM GLM2. In the dmPFC in Study 1, the performance of both models was equivalent.

| Contrast & Region | x | y | z | Cluster size | $T_{S1}$ | $T_{S2}$ |
|---|---|---|---|---|---|---|
| ***Arbitration signal (emulation reliability)*** | | | | | | |
| ACC | 0 | 39 | 3 | 155 | 4.34 | 4.14 |
| Right vlPFC / insula | 53 | 32 | 1 | 376 | 3.76 | 3.71 |
| Right mid/sup temporal | 48 | -21 | -7 | 61 | 3.84 | 3.67 |
| Left postcentral / supramarginal | -58 | -29 | 18 | 207 | 3.49 | 3.99 |
| Right supramarginal / inferior parietal | 65 | -31 | 26 | 185 | 3.65 | 3.67 |
| Left fusiform gyrus | -25 | -71 | -15 | 146 | 3.77 | 3.50 |
| Right fusiform gyrus | 30 | -69 | -10 | 69 | 3.67 | 3.42 |
| dACC | 5 | 17 | 31 | 31 | 3.71 | 3.42 |
| Mid-cingulate cortex | 15 | -21 | 41 | 58 | 3.29 | 3.42 |
| Left insula | -40 | -9 | -7 | 177 | 3.72 | 3.41 |
| Left anterior insula | -43 | 12 | -12 | 34 | 3.11 | 3.39 |
| SMA / preSMA | 8 | -9 | 76 | 143 | 3.36 | 3.67 |
| Left pSTS/TPJ | -58 | -54 | 13 | 68 | 3.05 | 3.14 |
| ***Emulation learning signal (token KL divergence) during feedback*** | | | | | | |
| Left anterior insula | -35 | 17 | -7 | 117 | 4.50 | 4.17 |
| Right anterior insula | 35 | 19 | -10 | 179 | 3.81 | 4.25 |
| Right IFG | 43 | 9 | 26 | 328 | 3.64 | 3.33 |
| Left IFG | -40 | 7 | 28 | 171 | 3.37 | 3.33 |
| Right caudate / thalamus | 8 | -1 | 8 | 106 | 3.95 | 3.69 |
| Left fusiform gyrus | -35 | -56 | -15 | 37 | 4.13 | 3.87 |
| Right inferior occipital | 30 | -86 | -10 | 54 | 3.31 | 3.80 |
| Left inf-sup parietal / precuneus | -25 | -71 | 38 | 389 | 3.60 | 4.47 |
| Right superior occipital / parietal | 28 | -61 | 41 | 38 | 3.28 | 3.73 |
| Right inferior parietal | 50 | -34 | 46 | 187 | 3.32 | 3.27 |
| Right occipital / cuneus | 18 | -76 | 6 | 345 | 3.41 | 3.47 |
| Left mid-sup frontal / precentral | -23 | 2 | 51 | 169 | 3.30 | 3.38 |
| Right mid-sup frontal / precentral | 30 | 9 | 58 | 136 | 3.34 | 3.18 |
| SMA / preSMA | 5 | 22 | 48 | 351 | 3.80 | 3.77 |
| Right TPJ / pSTS | 55 | -44 | 23 | 44 | 3.28 | 3.22 |

| | | | | | | |
|---|---|---|---|---|---|---|
| ***Imitation learning signal (action change) during feedback*** | | | | | | |
| SMA / preSMA | -5 | 4 | 66 | 47 | 3.21 | 3.11 |
| Left inferior parietal | -38 | -54 | 41 | 318 | 3.66 | 3.65 |
| Right inferior parietal | 50 | -39 | 48 | 129 | 3.63 | 3.55 |
| Left dlPFC | -45 | 32 | 31 | 53 | 3.46 | 3.59 |
| Left anterior insula | -35 | 22 | -10 | 45 | 3.47 | 3.43 |
| Left mid-sup frontal / precentral | -18 | 14 | 63 | 119 | 3.23 | 3.18 |
| Left IFG | -45 | 4 | 26 | 35 | 2.95 | 3.38 |
| Precuneus | 20 | -69 | 61 | 34 | 2.75 | 2.74 |
| ***Previous action unavailable > available during slot machine presentation*** | | | | | | |
| SMA / preSMA | -8 | 12 | 51 | 1132 | 5.18 | 5.17 |
| Left anterior insula | -35 | 22 | -2 | 237 | 5.35 | 5.08 |
| Right anterior insula | 38 | 24 | -2 | 253 | 4.89 | 4.63 |
| Left IFG | -45 | 4 | 28 | 531 | 4.97 | 5.04 |
| Right IFG | 45 | 7 | 26 | 270 | 4.38 | 4.46 |
| Left sup occipital / inf-sup parietal | -33 | -84 | 28 | 1077 | 4.22 | 4.21 |
| Right inf-sup parietal / occipital | 53 | -36 | 48 | 1505 | 4.45 | 4.52 |
| Right mid-sup frontal / precentral | 28 | -6 | 53 | 408 | 4.12 | 4.18 |
| Right inferior temporal | 50 | -56 | -10 | 153 | 4.20 | 3.98 |
| Left occipital / cuneus | -15 | -81 | 3 | 58 | 3.82 | 3.06 |
| Right occipital / cuneus | 15 | -69 | 13 | 50 | 4.31 | 3.31 |
| Left caudate | -10 | 2 | 3 | 47 | 3.45 | 3.44 |
| Right caudate | 15 | 12 | 1 | 54 | 3.97 | 3.36 |
| Left inferior parietal | -50 | -41 | 43 | 54 | 3.25 | 4.08 |
| Right cuneus / precuneus | 23 | -59 | 21 | 46 | 3.31 | 3.48 |
| Right dlPFC | 48 | 32 | 26 | 60 | 3.29 | 3.63 |
| Left inferior occipital/temporal / fusiform | -38 | -71 | -12 | 168 | 3.33 | 3.71 |
| Right fusiform gyrus | 30 | -71 | -12 | 32 | 3.37 | 3.17 |
| ***Token value during token presentation*** | | | | | | |
| ACC | -8 | 39 | -2 | 58 | 4.26 | 3.78 |
| mOFC / vmPFC | -10 | 64 | 1 | 30 | 3.02 | 2.89 |
| ***Emulation choice probability during self-choice (p<0.001 unc)*** | | | | | | |
| *mOFC / vmPFC* | *-8* | *59* | *-10* | *210* | 2.75 | 2.96 |
| ***Negative Imitation choice probability during self-choice (p<0.001 unc)*** | | | | | | |
| *Right anterior insula* | *38* | *29* | *-2* | *67* | 2.87 | 3.46 |
| *dmPFC / preSMA / SMA* | *3* | *24* | *43* | *64* | 2.59 | 2.63 |

**Table S4. SPM GLM3 conjunction analyses results.** Conjunction maps between the second-level T-maps of studies 1 and 2 were thresholded at $P_{conjunction}<0.0001$ uncorrected, followed by whole-brain cluster level family-wise error correction at $P_{FWE}<0.05$ (equivalent to cluster size k≥30 contiguous voxels). For the last two contrasts, emulation choice probability and negative imitation choice probability, no cluster survived the above threshold so a slightly more lenient cluster-forming threshold of $P_{conjunction}<0.001$ uncorrected was used for exploratory purposes. x, y, z represent MNI coordinates. $T_{S1}$ and $T_{S2}$ denote the T-value for Study 1 and Study 2, respectively. mOFC: medial orbitofrontal cortex. vmPFC: ventromedial prefrontal cortex. ACC: anterior cingulate cortex. IFG: inferior frontal gyrus. SMA: supplementary motor area. dmPFC: dorsomedial prefrontal cortex. dlPFC: dorsolateral prefrontal cortex. TPJ: temporoparietal junction. pSTS: posterior superior temporal sulcus.

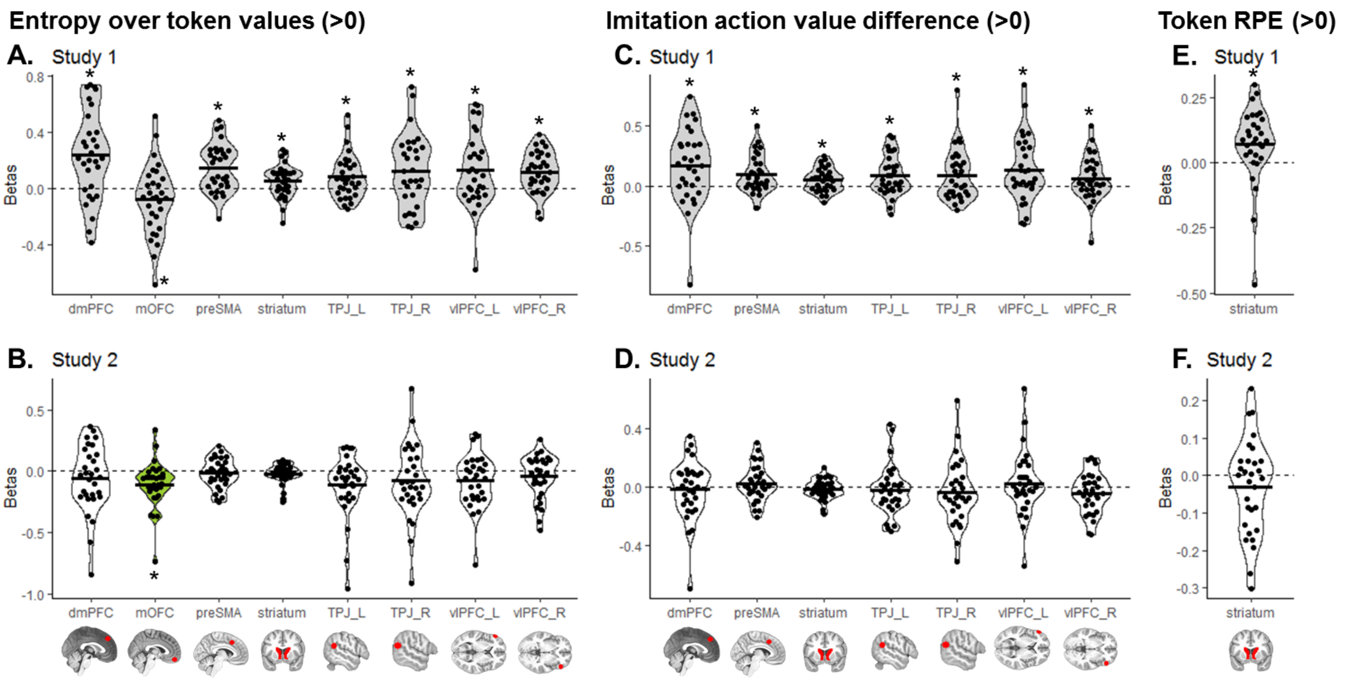


**Figure S1. Pre-registered ROI results – additional contrasts.** For the remaining contrasts in SPM GLM1 and GLM2 not shown on main text **Figures 4-6**, mean signal was extracted from each pre-registered ROI. Regions with significant signals in Study 1, plotted in grey, were selected as hypotheses and a priori ROI for Study 2. Green plots represent significant effects in Study 2, confirming the a priori hypothesis from Study 1. White plots represent hypotheses that were not confirmed in Study 2. Dots represent individual subjects and the black bar represents the mean beta value for each regressor. T-tests: * P<0.05. The same results were found using non-parametric permutation tests.

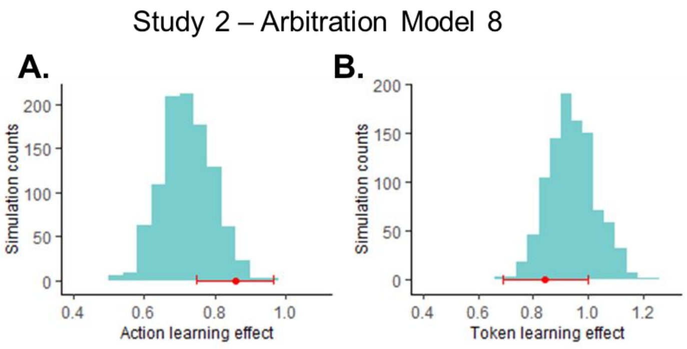


**Figure S2. Action and token learning effects are also captured by arbitration model 8 in Study 2.** We tested how well Arbitration Model 8, which performed best in Study 2 based on BIC values, was able to capture the two behavioral effects obtained by a simple logistic regression: the action learning effect (**A**) and the token learning effect (**B**). The red data point above the X-axis depicts the true effect from the data, and the histogram shows the distribution of the recovered effects from the model-generated data.

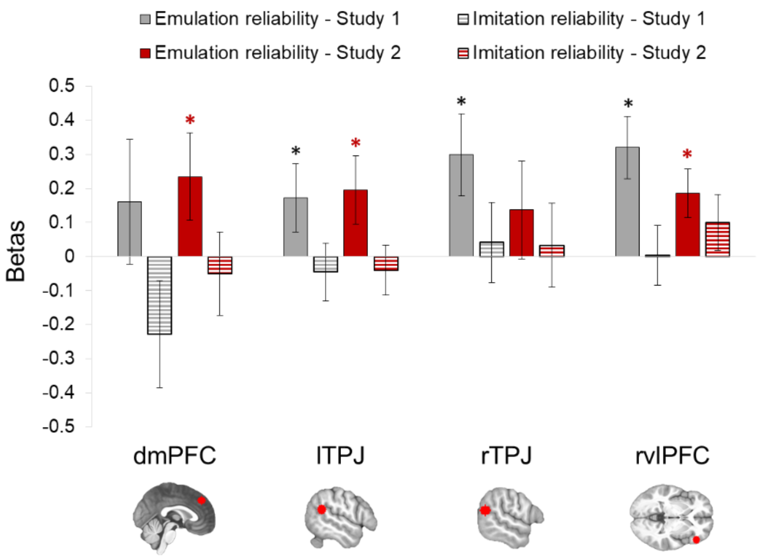


**Figure S3. Reliability difference signals are only driven by positive tracking of emulation reliability.** In Study 1, significant reliability difference signal was found in four ROIs: dmPFC, left and right TPJ, and right vlPFC. We extracted emulation (solid fill) and imitation (horizontal stripes fill) reliability signals separately in each of these ROIs and for each study (Study 1: grey, Study 2: red), to test whether the reliability difference signal is driven by both positive tracking of emulation reliability and negative tracking of imitation reliability. However, we find that this was not the case, instead only emulation reliability was found to be significant represented in the ROIs. Error bars represent SEM. T-tests: * P<0.05. The same results were found using non-parametric permutation tests.

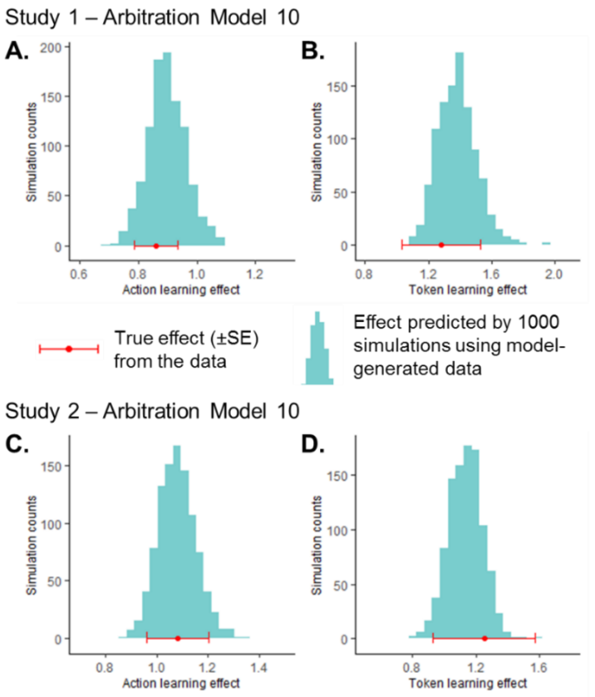


**Figure S4. Action and token learning effects are captured by arbitration model 10.** We tested how well Arbitration Model 10, which arbitrates between the original emulation strategy and a simpler 1-step imitation strategy, was able to capture the two behavioral effects obtained by a simple logistic regression: the action learning effect (**A, C**) and the token learning effect (**B, D**), in both Study 1 (top) and Study 2 (bottom). The red data point above the X-axis depicts the true effect from the data, and the histogram shows the distribution of the recovered effects from the model-generated data.

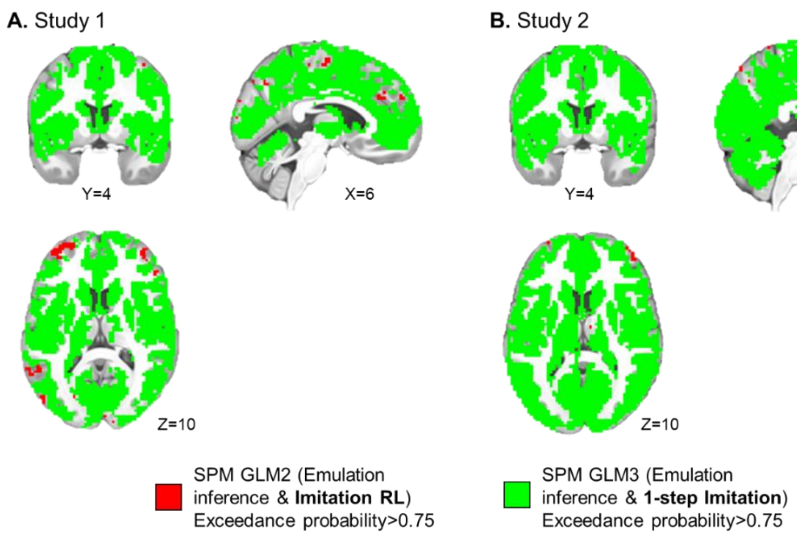


**Figure S5. fMRI Bayesian Model Selection results.** The maps shown represent grey matter voxels with exceedance probabilities greater than 0.75, in red for SPM GLM2 (representation of emulation inference signals and imitation RL signals) and in green for SPM GLM3 (representation of emulation inference signals and 1-step imitation signals). In both Study 1 (**A**) and Study 2 (**B**), SPM GLM3 was found to provide a better account of variations in the BOLD signal in a vast majority of grey matter voxels.

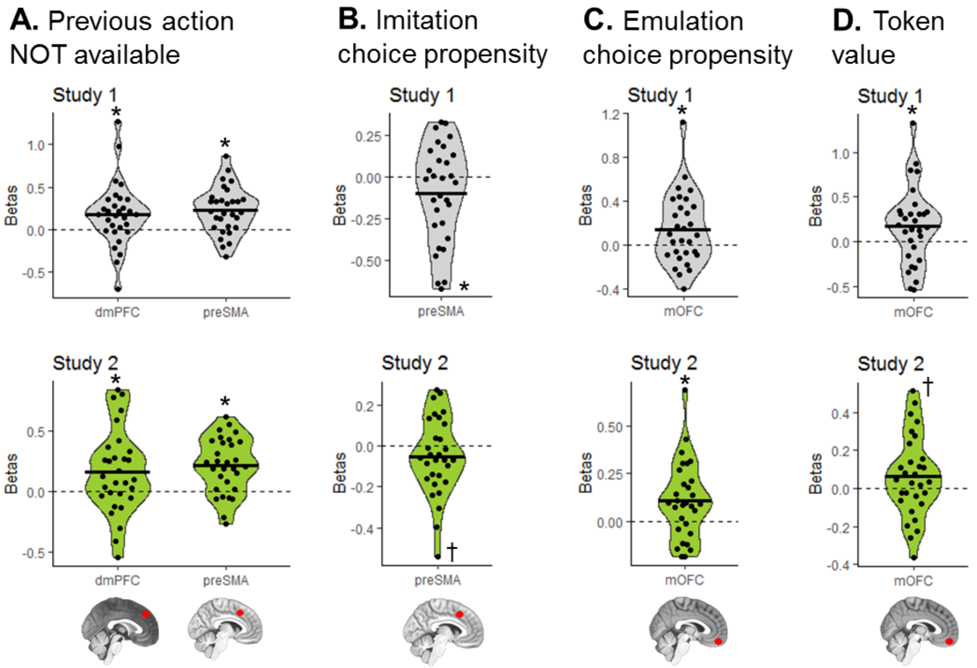


**Figure S6. SPM GLM3 additional contrasts ROI results.** Mean signal was extracted from pre-registered ROIs for each remaining contrast of interest in SPM GLM3 not shown on main text **Figures 5-6**: (**A**) whether the partner's previous action is unavailable (vs available) on the current trial, (**B-C**) the probability to choose according to imitation (**B**) or emulation (**C**) at the time of self-choice, and (**D**) the value of the token shown on screen during token presentation. Regions with significant signals in Study 1 (top panels), plotted in grey, were selected as hypotheses and a priori ROI for Study 2 (bottom panels). Dots represent individual subjects and the black bar represents the mean beta value for each regressor. T-tests: * $P<0.05$, † $P\leq0.07$. The same results were found using non-parametric permutation tests.
